# Innate sensitivity and plastic mechanisms in auditory cortex for reliable maternal behavior

**DOI:** 10.1101/2020.03.11.987941

**Authors:** Jennifer K. Schiavo, Silvana Valtcheva, Chloe Bair-Marshall, Soomin C. Song, Kathleen A. Martin, Robert C. Froemke

**Affiliations:** Skirball Institute for Biomolecular Medicine, New York University School of Medicine, New York, NY, USA; Neuroscience Institute, New York University School of Medicine, New York, NY, USA; Department of Otolaryngology, New York University School of Medicine, New York, NY, USA; Department of Neuroscience and Physiology, New York University School of Medicine, New York, NY, USA; Center for Neural Science, New York University, New York, NY, USA; Howard Hughes Medical Institute Faculty Scholar

## Abstract

Infant cries evoke powerful responses in parents^1–4^. To what extent are parental animals innately sensitive to neonatal vocalizations, or might instead learn about key vocal cues for appropriate parenting responses? In mice, naive virgins do not recognize the meaning of pup distress calls, but begin to retrieve pups to the nest following cohousing with a mother and litter^5–8^. These isolation calls can be variable, requiring co-caring virgins to generalize across features for reliable retrieval^9, 10^. Here, using behavioral studies combined with two-photon imaging and whole-cell recordings, we show that the onset of maternal behavior in mice results from the interaction between innate sensitivities and experience-dependent processes. We found that pup calls with inter-syllable intervals (ISIs) ranging from 75 to 375 ms elicited pup retrieval, and experienced auditory cortex generalized across these ISIs. In contrast, naive cortex was narrowly tuned to the most common or ‘prototypical’ ISIs due to enhanced short-term depression of inhibitory inputs. Behavioral testing revealed that naive virgins were also more sensitive to prototypical calls than calls at other rates. Inhibitory and excitatory spiking and synaptic responses were initially mismatched in naive cortex, with untuned inhibition and overly-narrow excitation. Monitoring neuronal populations over cohousing revealed that excitatory neuronal responses broadened to represent a wide range of ISIs, while inhibitory neurons sharpened to form a perceptual boundary. Finally, we presented synthetic calls during cohousing and observed that neural and behavioral responses adjusted to match these statistics. Using inhibitory optogenetics, we found that auditory cortical activity was required to learn about specific features, whereas the oxytocinergic system was generally recruited for retrieval learning and plasticity in temporal tuning. Neuroplastic mechanisms therefore build on an innate sensitivity in the auditory cortex, enabling rapid plasticity for reliable parenting behavior.

---

Parents must quickly respond to neonatal vocalizations signaling physiological needs^1–4^. Mammalian infants produce various types of vocalizations such as laughing, babbling, or cries, which have distinct spectro-temporal features defining their behavioral meaning. However, appropriate behavioral responses are particularly difficult when vocalizations with the same meaning vary due to inter- and intra-individual differences in vocal production^2, 4, 11^. This requires listeners to generalize across acoustic features to recognize the behavioral relevance and act appropriately^12–14^. While vocal perception is typically learned^15^, aspects of parental care might also be hard-wired within relevant brain areas (or biased from pre-parental experience) given the biological imperative to ensure offspring survival. Here we ask to what degree the auditory cortex is intrinsically tuned to particular vocal features prior to parental experience, and what neuroplastic mechanisms underlie auditory learning for maternal behavior.

Rodent mothers (‘dams’) retrieve lost pups back to the nest based on ultrasonic distress calls^7–9^. These isolation calls contain spectro-temporal qualities distinct from adult vocalizations and other pup vocalizations (e.g., wriggling calls eliciting nursing), and are organized into temporally-modulated bouts around 3-8 Hz with syllables in the ultrasonic range (∼50-80 kHz)^9, 10, 16^. Distress calls vary within and across individuals, but the purpose of these calls is the same: to increase arousal and elicit retrieval behavior. This requires caretakers to generalize across natural variability in key vocal features for rapid pup retrieval. Pup-naive virgin mice do not initially behaviorally respond to these isolation calls, but can begin to retrieve following experience with pups (‘experienced virgins’)^5–8^. The emergence of alloparenting in virgins thus allows us to dissect the innate vs. learned components of pup call recognition for the onset of reliable maternal behavior.

We first wondered whether syllable repetition rate is a key feature driving pup retrieval. The temporal structure of isolation calls is effective for recognition in noisy environments^16^, and other pup vocalizations do not occur in rhythmic bouts^17^. We recorded calls emitted from isolated pups (n=32 ∼3 min. recordings; see methods), and measured the probability of observing various inter-syllable intervals (n=4,540 ISIs, **Fig. 1a**). We selected calls containing ISIs near the median of the distribution to serve as ‘prototypical’ pup calls (ISI:150-200 ms, ‘175 ms’). To generate a library of calls with various repetition rates, we morphed prototypical calls by adding or subtracting time between syllables (i.e., increasing or decreasing ISIs), ensuring prototypes and their longer or shorter morphs were matched in all other features (**Fig. 1a, Extended Data Fig. 1**).

**Figure 1.**
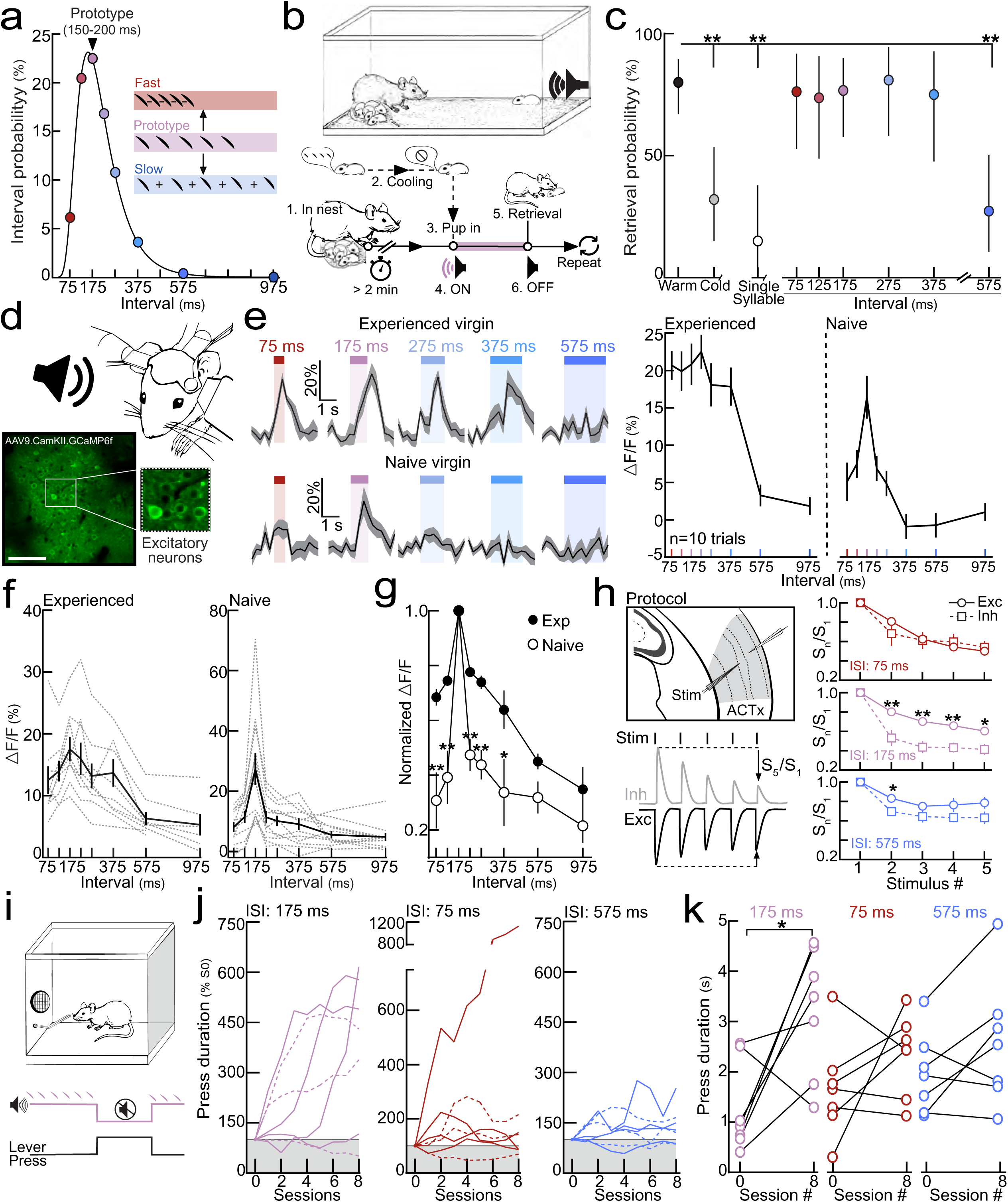
Temporal statistics drive behavioral and cortical responses to pup calls in naive and experienced virgins. **a**, Lognormal probability distribution of ISIs (each dot represents binned ISIs centered at each point ±25 ms). Prototypes were morphed by adding or subtracting time between syllables. Colors denote ISI bins throughout the manuscript. **b,** Experimental protocol for dubbing over cold pups. **c,** Retrieval of ‘dubbed over’ cold pups. Females retrieved vocalizing pups and cold pups dubbed over with morphs with ISIs between 75 and 375 ms at similar rates (75:375 ms, p>0.05; Binomial test). Error bars, 95% binomial CI. **d,** In vivo two-photon Ca^2+^ imaging. Scale bar, 75 μm. **e**, Example ΔF/F Ca^2+^ transients (left) and quantification (right) from sample cells imaged in an experienced and naive virgin. **f,** Excitatory neuronal tuning across experienced and naive virgins. **g,** Tuning normalized to prototypes. Responses were more similar between prototypes and ISIs 75:375 ms (higher normalized ΔF/Fs) in experienced virgins (75:375 ms, p<0.05; Student’s unpaired one-tailed t-test). **h,** Left, protocol for in vitro whole-cell recordings in virgin auditory cortex. Right, IPSCs depressed faster than EPSCs at prototypical repetition rates (Stim 2:5, p<0.01). **i**, Operant test: a lever press turns off pup calls for the duration of the press. **j**, Learning trajectories (normalized to baseline). **k**, Virgins listening to prototypes, but not fast or slow morphs, significantly increased lever press duration by session 8 (p=0.03, Student’s paired two-tailed t-test).

We then devised a behavioral assay to test which repetition rates elicited retrieval in experienced females. Pups were anesthetized on ice and placed under a speaker in the behavioral arena, allowing us to ‘dub over’ the cold pup with prototypical calls or spectrally-matched morphs (**Fig. 1b**). Experienced females retrieved cold, silent pups (n=8/25 trials) on significantly fewer trials than warm, vocalizing pups (n=44/55, p<0.0001). Females retrieved cold pups dubbed over with prototypical calls (n=23/30), as well as temporal morphs as long as ISIs fell between 75 and 375 ms (n=59/77). ISIs slower than 575 ms (n=6/22) and single syllables separated by 30 s (n=3/20) were not effective in eliciting retrieval (p<0.0001), suggesting a series of syllables in close succession was required (**Fig. 1c, Extended Data Fig. 2**). Notably, ISIs slower than ∼375 ms rarely occurred in our distribution (<1%), suggesting females may preferentially learn about prevalent vocal features. We observed similar results in a y-maze paradigm in which mice were tested for approach towards speakers playing prototypical calls versus spectrally-matched temporal morphs. Experienced females approached speakers playing prototypes significantly more as competing ISIs became slower (575:1000 ms, p=0.005; **Extended Data Fig. 3**). Taken together, these data suggest repetition rate is a critical feature for call recognition.

Next, we determined how maternal experience shapes the encoding of pup calls across temporal statistics in the auditory cortex. The left auditory cortex is a critical brain region for pup retrieval^8, 18^, and experience-dependent processing of call repetition rates has been previously reported in this brain region^19^. We performed two-photon calcium imaging of layer 2/3 excitatory neurons in the left auditory cortex of awake, head-fixed experienced and naive virgins^20^ (**Fig. 1d; Extended Data Fig. 4**). Robust responses were observed to prototypical calls in ∼16% of layer 2/3 excitatory neurons regardless of maternal experience (Experienced=15.8%, Naive=17.9%; **Extended Data Fig. 4a**). In naive virgins, excitatory neurons responsive to prototypes had reduced responses to the same call morphed in the temporal domain. In contrast, prototype-responsive neurons in experienced virgins had substantial responses across all ISIs until ∼575 ms, where responses were reduced (Sample cell, **Fig. 1e;** Sample population, **Extended Data Fig. 4b**). These temporal tuning profiles were consistent across virgins (Experienced, N=9 mice; Naive, N=12; **Fig. 1f**). Call-evoked ΔF/Fs in experienced cortex were also correlated with the probability of observing a given ISI, indicating neurons preferentially responded to high probability statistics (Pearson’s r=0.84, p=0.009; **Extended Data Fig. 4c**). Robust responses across ISIs were left lateralized in experienced virgins (75:375 ms, p=0.003; **Extended Data Fig. 5**), consistent with the specialized role the left temporal lobe plays in processing vocalizations^8, 9, 21, 22^.

In order to compare large populations of cortical neurons, we normalized neuronal responses to each cell’s response to the prototype. Higher values were indicative of a more similar response between a given prototype and the spectrally-matched morph (‘normalized ΔF/F’, **Fig. 1g**). Temporal tuning in experienced cortex was broader than in naive virgins, resulting in significantly higher normalized ΔF/Fs across the range of ISIs that elicited retrieval of ‘dubbed over’ cold pups (75:375 ms, p<0.05; **Fig. 1c,g**). This broad temporal tuning in experienced cortex may enable generalization across calls for reliable retrieval, as temporal tuning in females retrieving on 10-30% of trials was significantly sharper prior to the onset of robust retrieval behavior (p<0.0001; **Extended Data Fig. 6**). Narrow temporal tuning in naive cortex, however, may reflect an intrinsic sensitivity for vocalizations containing prototypical features. Interestingly, narrowly-tuned responses in naive virgins were specific to pup calls. There was no difference tuning across temporally-modulated pure tones in naive and experienced females (**Extended Data Fig. 7a-c**), and neurons were more broadly tuned to temporally-modulated tones than calls in naive virgin cortex (all ISIs, p<0.05; **Extended Data Fig. 7d,e**).

To determine whether synaptic mechanisms could confer this hardwired sensitivity to prototypical pup calls, we performed in vitro voltage-clamp recordings in pup-naive auditory cortical slices. Specifically, we measured excitatory (EPSC) and inhibitory (IPSC) postsynaptic currents following repeated pulses of extracellular stimulation (**Fig. 1h**-left). At fast (ISI:75 ms, ∼13 Hz) and slow (ISI:575 ms, ∼1.7 Hz) stimulation rates, EPSCs and IPSCs were similarly depressed (n=8-13 cells). However, IPSCs adapted out significantly faster than EPSCs at prototypical repetition rates of ∼5-6 Hz (n=11-14 cells, p<0.01 for stim 2:5; **Fig. 1h**-right). In this way, inhibitory adaptation could provide a synaptic basis for intrinsic sensitivity to prototypical calls in naive cortex. These dynamics could also enable and/or result from developmental plasticity, consistent with a previous study that found heightened plasticity in the rat auditory cortex for stimuli presented at 6 Hz^23^.

Could this prewired sensitivity for prototypical calls serve to speed up learning during early maternal experience? To test the hypothesis that prototypical calls are particularly salient to naive virgins, we trained virgin mice to press a lever to turn off continuously playing prototypical calls (175 ms), fast morphs (75 ms), or slow morphs (575 ms) for the duration of the lever press (N=7 mice/group; **Fig. 1i**, see methods). Naive virgins learned to turn off prototypical calls via lever press over eight days (**Fig. 1j**, pink), and lever press duration significantly increased by session 8 (p=0.03; **Fig. 1k**). No change in lever press duration was observed for virgins in the fast (p=0.24) or slow (p=0.13) morph groups (**Fig. 1k**). We did not find any significant change in the number of lever presses per session in any group (**Extended Data Fig. 8**). These data suggest naive virgins find pup calls with prototypical ISIs particularly salient, or even aversive, as they were primarily motivated to turn off prototypes. This intrinsic sensitivity of female left auditory cortex resulting from excitatory-inhibitory dynamics might lead to enhanced arousal and accelerate the onset of maternal learning as virgins are motivated to placate vocalizing pups.

Next, we wondered whether inhibitory neurons are also differentially tuned to pup call statistics in naive and experienced females. We selectively expressed GCaMP6f in inhibitory neurons using a cell-type specific virus^24^ (**Fig. 2a**). Inhibitory neurons appeared broadly tuned in experienced and naive virgins (**Fig. 2b**), resulting in similar tuning profiles across animals (N=5-6 mice, p>0.05 for all ISIs; **Fig. 2c, Extended Data Fig. 9**). However, there was one key difference in inhibitory neuronal tuning; the slope of the tuning curve at the transition between ISIs that elicited retrieval (75:375 ms) and those that did not (575:975 ms; **Fig. 2b,c**, blue shading) was significantly sharper in experienced cortex (p=0.03; **Fig. 2d**). We hypothesize this sharp transition could enhance discrimination between salient and non-salient call features in retrieving females^25^. On the other hand, a shallow slope in naive cortex indicates that, although inhibitory neurons are broadly tuned^26^, this population is still relatively untuned to relevant statistics. Furthermore, excitatory and inhibitory neuronal responses were matched in experienced cortex, responding broadly across the range of behaviorally-salient ISIs (ISIs 75:375; **Fig. 2e**). This is in contrast to the naive cortex, in which inhibitory neurons were more broadly tuned than excitatory neurons (For all ISIs, p<0.001; **Fig. 2f**).

**Figure 2.**
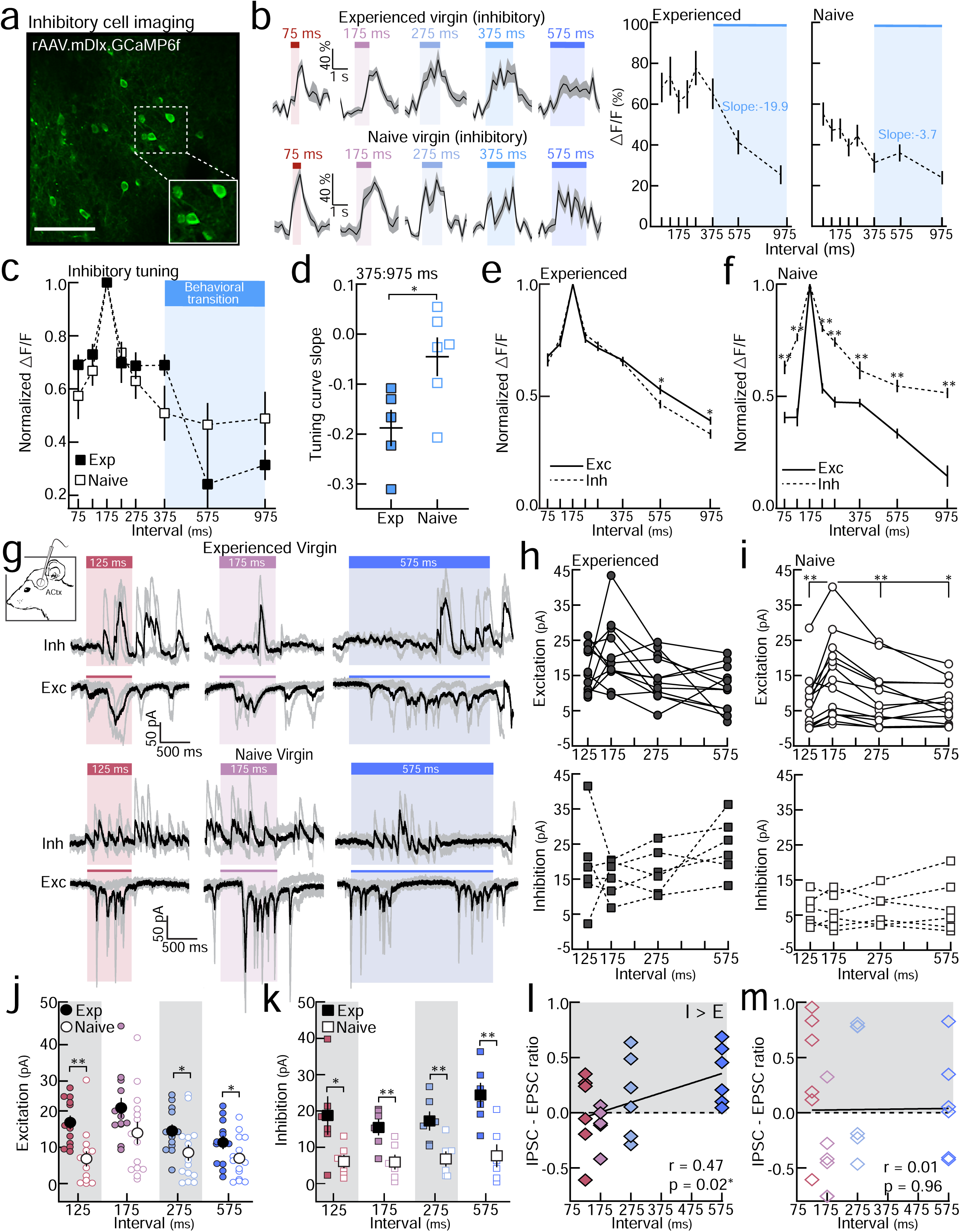
Excitatory and inhibitory tuning and synaptic responses are altered by maternal experience. **a**, GCaMP6f in interneurons. Scale bar, 75 μm. **b**, Example ΔF/F Ca^2+^ transients (left) and quantification (right) from inhibitory neurons. Blue shading denotes behavioral transition from Fig. 1c. **c**, Inhibitory temporal tuning (norm. to prototype) across experienced and naive virgins (All ISIs, p>0.05). **d,** Slope of tuning curves (375:975 ms) (p=0.03). **e,f,** Excitatory (n=366 single-cell tuning curves) and inhibitory (n=260) neuronal tuning was similar in experienced cortex for central ISIs (575 ms, p=0.02; 975 ms, p=0.03), but mismatched in naive cortex across all ISIs (Exc (n=268) vs. Inh (n=128): **p<0.001). **g**, Example in vivo voltage-clamp recordings from auditory cortical neurons. **h,** Synaptic tuning in experienced cortex. For all ISIs, p>0.05 (repeated measures one-way ANOVA with Bonferroni correction). PSC (pA)=Area (pA*ms) divided by stim. duration (ms). **i,** Synaptic tuning in naive cortex. EPSCs (top): prototype vs. all ISIs: p<0.05. IPSCs: prototype vs. all ISIs, p>0.99 (repeated measures one-way ANOVA with Bonferroni correction). **j,** EPSCs evoked by non-prototypical ISIs were larger in experienced cortex (125, p=0.0008, 275, p=0.03, 575, p=0.04; Student’s unpaired one-tailed t-test). **k,** Evoked IPSCs were larger in experienced cortex (p<0.05). **l,m,** IPSC-EPSC ratio was positively correlated with ISI duration in experienced cortex.

These imaging data demonstrate that maternal experience adjusts the relative balance of excitatory and inhibitory tuning at the output level. To directly determine how experience-dependent temporal tuning arises from synaptic inputs, we performed in vivo voltage-clamp recordings from layer 2/3 auditory cortical neurons in isoflurane-anesthetized experienced and naive females^8^ (**Fig. 2g**). In experienced females, prototype- and morph-evoked EPSCs did not significantly differ (n=12 neurons; **Fig. 2h**-top), and were negatively correlated with ISI duration (Pearson’s r=-0.37, p=0.01; **Extended Data Fig. 10a**). In naive cortex, evoked excitation was substantially weaker in response to morphs (n=14, p<0.01 for all ISIs; **Fig. 2i**-top), and EPSCs were uncorrelated with ISI duration (Pearson’s r=-0.13, p=0.35; **Extended Data Fig. 10c**). To test the hypothesis that maternal experience enhances excitatory drive across relevant statistics, we directly compared EPSC magnitudes in naive and experienced cortex. While prototype-evoked EPSCs were similar between groups (p=0.05), responses to non-prototypical ISIs were enhanced in experienced females (p<0.05; **Fig. 2j**). In contrast, prototype- and morph-evoked IPSCs did not significantly differ within a neuron (**Fig. 2h,i-**bottom) and did not track ISI duration in either group (n=6 neurons/group; **Extended Data Figure 10b,d**). IPSCs were significantly larger in experienced cortex overall (p<0.05; **Fig. 2k**), as would be expected to balance the increase in excitatory drive that we observed following maternal experience^8^.

Most importantly, these results suggest that inhibitory inputs in experienced cortex may define the boundary for relevant call statistics. In prototype-responsive neurons in experienced cortex, EPSCs evoked by 575 ms morphs did not significantly differ from the prototype (p=0.18; **Fig. 2h**-top), an interval where output tuning was diminished (**Fig. 1e-g, Fig. 2c**). However, when we calculated the relative difference in excitatory-inhibitory magnitudes within a neuron (n=6 neurons) (‘IPSC-EPSC ratio’: 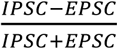), we found this ratio to be positively correlated with ISI duration (Pearson’s r=0.47, p=0.02; **Fig. 2l**). This suggests that inhibition is higher than excitation in response to slow morphs in experienced females, which could sharpen temporal tuning at the transition between salient and non-salient ISIs. No such correlation was present in naive cortex (n=6, Pearson’s r=0.01, p=0.96; **Fig. 2m**). Thus, while inhibitory neurons form a sharp boundary to slow ISIs following maternal experience (**Fig. 2b-d**), net postsynaptic inhibitory impact on cortical neurons is enhanced. This would allow inhibitory cell firing to accurately track statistics of perceived pup calls, while individual downstream neurons adjust synaptic strengths depending on the specific computations these cells perform for pup call detection. Taken together, these imaging and electrophysiological results demonstrate that the naive auditory cortex is narrowly tuned to prototypical calls, partially resulting from weak and untuned inhibition in conjunction with reduced excitatory drive to non-prototypical ISIs. Maternal experience results in enhanced excitatory and inhibitory inputs, broadening the temporal tuning of auditory cortical neurons.

When does this plasticity in temporal tuning occur during early maternal experience? We expressed GCaMP6f in either excitatory or inhibitory neurons, and monitored neuronal populations (and in some cases, the same cells) over several days of cohousing with a dam and litter. Virgins were tested for retrieval every 12 hours and imaged at least every 24 hours, including imaging after the first pup retrieval and 24 hours later (**Fig. 3a**). Spatial cross-correlations of surrounding pixels were used to identify the same neurons over days^27^ (**Fig. 3b,c**; see methods). At the single-cell level, we observed substantial differences in the stability of prototype-responsive neurons (**Fig. 3d**). While inhibitory neuronal responses to prototypes were correlated before and after retrieval onset (n=93 neurons, Pearson’s r=0.47, p<0.0001), distinct excitatory neurons were recruited following maternal experience (n=116, Pearson’s r=0.16, p=0.08; **Fig. 3e**). Of the neurons that were prototype-responsive at baseline and successfully tracked, ∼18.1% of excitatory neurons (n=21/116 cells) and ∼71.0% of inhibitory neurons (n=66/93) remained responsive in the final imaging session.

**Figure 3.**
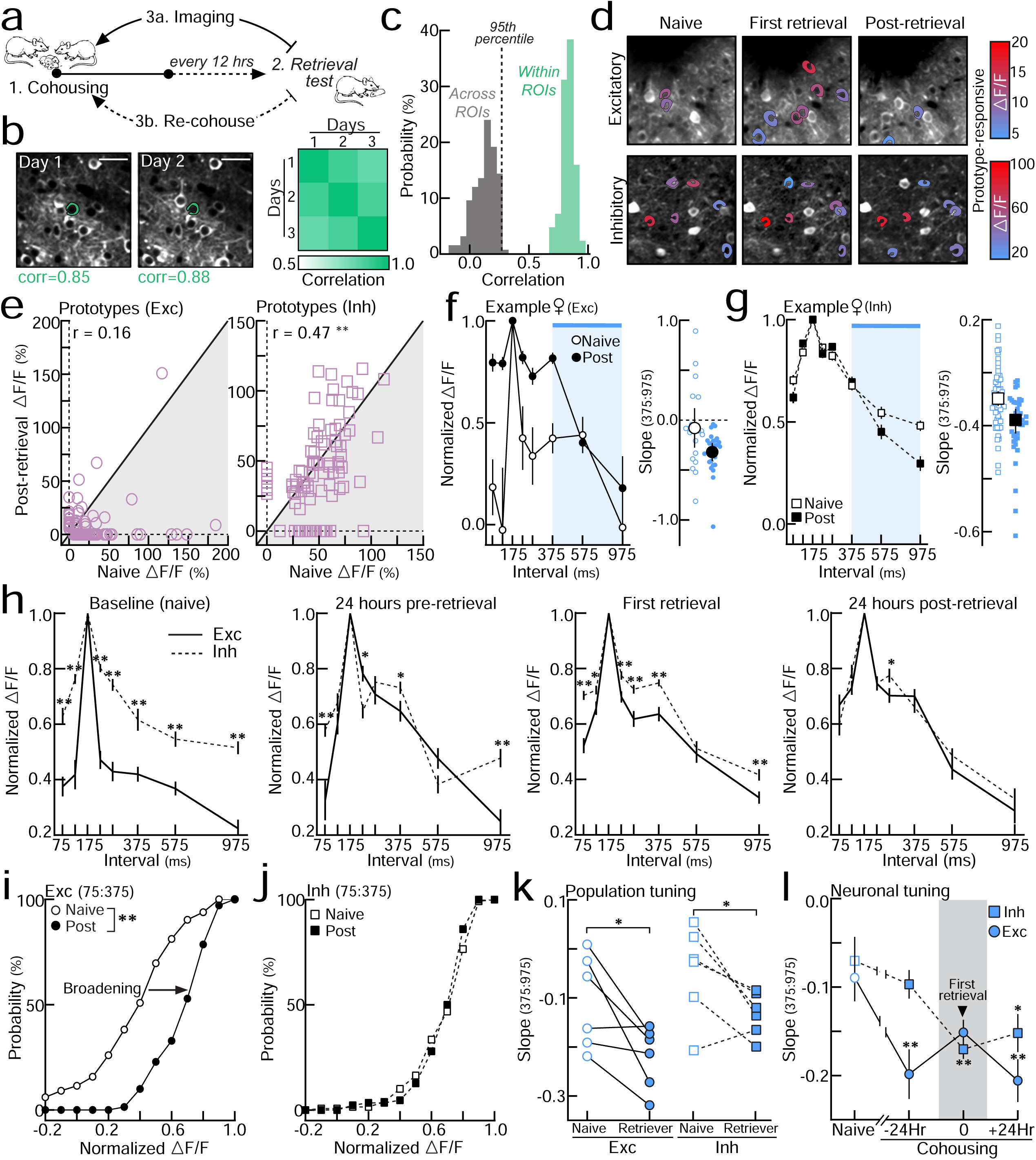
Cohousing with pups results in coordinated plasticity of excitatory and inhibitory neuronal tuning. **a,** Protocol for chronic two-photon imaging and retrieval testing. **b,c,** Spatial cross-correlations of pixels surrounding an ROI for single-cell identification (see methods). **d,** Tracking of excitatory (top) and inhibitory (bottom) neurons. Colored cell bodies denote prototype-responsive cells based on ΔF/F. **e,** Correlations of prototype-evoked responses before and after retrieval. Zeros denote non-responsive cells. **f,g,** Example excitatory **(f)** and inhibitory **(g)** population tuning with a breakdown of each single-cell tuning curve slope at 375:975 ms. Blue shading denotes behavioral transition as in **2c**. **h,** Excitatory (n=165 single-cell tuning curves) and inhibitory (n=128) tuning was initially mismatched (**p<0.0001), but came to match 24 hours after first retrieval (Exc, n=70; Inh, n=86). Binned ±12 hours. **i,j**, Cumulative distributions of normalized ΔF/Fs averaged across 75:375 ms. Excitatory neurons **(i)** broadened tuning (Naive: n=165 single-cell tuning curves, post-retrieval: n=70; p<0.0001), while inhibitory tuning **(j)** did not change (Naive: n=128, post: n=86; p=0.16; Kolmogorov-Smirnov test). **k**,**l**, Excitatory and inhibitory tuning curves sharpened around 375:975 ms at the population (**k**, Exc, p=0.02; Inh, p=0.02; Student’s one-tailed paired t-test) and single-cell levels (**l, \***p<0.05, **p<0.01; one-way ANOVA with Bonferroni correction). Data aligned to first retrieval and binned ±12 hours.

Despite these single-cell dynamics, we identified consistent changes in excitatory and inhibitory temporal tuning at the population level during cohousing. Excitatory neuronal responses to ISIs ranging 75 to 375 ms became more similar to prototype-evoked responses (higher normalized ΔF/Fs), broadening single-cell tuning curves across behaviorally-salient ISIs (**Fig. 3f,h,i, Extended Data Fig. 11a**). This plasticity elaborated on the initial preference for prototypes, and also resulted in a sharpening of tuning curves between 375:975 ms at the population (N=6 mice, p=0.02; **Fig. 3k**) and single-cell levels (n=50-129 single-cell tuning curves, baseline vs. +24 hours: p=0.0005; **Fig. 3l**). This was different from what we observed for inhibitory neurons. Inhibitory cells did not broaden their tuning (**Fig. 3g,h,j, Extended Data Fig. 11b**), but rather sharpened selectively at the behavioral transition (375:975 ms) across animals (N=6 mice, p=0.02; **Fig. 3k**) and at the level of single-cell tuning curves (n=49-121 tuning curves, baseline vs. +24 hours: p=0.02; **Fig. 3l**). These opposing dynamics in inhibitory and excitatory neurons resulted in matched excitatory-inhibitory neuronal tuning to represent calls over a range of behaviorally-salient intervals. Interaction with pups was required for this plasticity, as non-cohoused controls only exposed to pup calls during consecutive imaging sessions showed no systematic changes in temporal tuning (**Extended Data Fig. 11c,d**).

Thus, two plastic processes occur with maternal experience: 1) enhanced neuronal responses across a range of behaviorally-relevant statistics, largely due to enduring enhancement of excitatory input; and 2) the emergence of a perceptual boundary in both excitatory and inhibitory populations. This indicates that there are multiple forms of plasticity acting in different directions. Excitatory cells respond more broadly to behaviorally-salient ISIs (75:375 ms), whereas the temporal tuning of inhibitory cells becomes sharper while maintaining enhanced postsynaptic inhibitory responses to rare call statistics (**Fig. 4a**). Does this re-tuning of cortical neurons reflect a pre-determined range of intervals, or can the relevant statistics be altered just by manipulating the acoustic environment during cohousing? To test the hypothesis that virgins generalize across call statistics heard during initial maternal experience, we cohoused naive virgins with a dam and litter in a cage containing a speaker over the nest. In addition to what calls might be experienced during cohousing, these virgins also heard slow morphs (ISI:575 ms) presented from the speaker every three hours in alternating 12-hour blocks (‘Cohoused+575’, **Fig. 4b**). We then used in vivo two-photon calcium imaging to assess neuronal temporal tuning 24 hours after retrieval onset. Compared to virgins with standard cohousing experience (‘Cohoused’, N=5 mice), prototype-responsive neurons in the ‘Cohoused+575’ group (N=6) were more broadly tuned with higher normalized ΔF/Fs in response to 575 ms morphs (575, p=0.009; **Fig. 4c, Extended Data Fig. 12a**). These data suggest we were able to expand temporal tuning curves to encompass 575 ms ISIs by simply manipulating the call statistics heard during cohousing.

**Figure 4.**
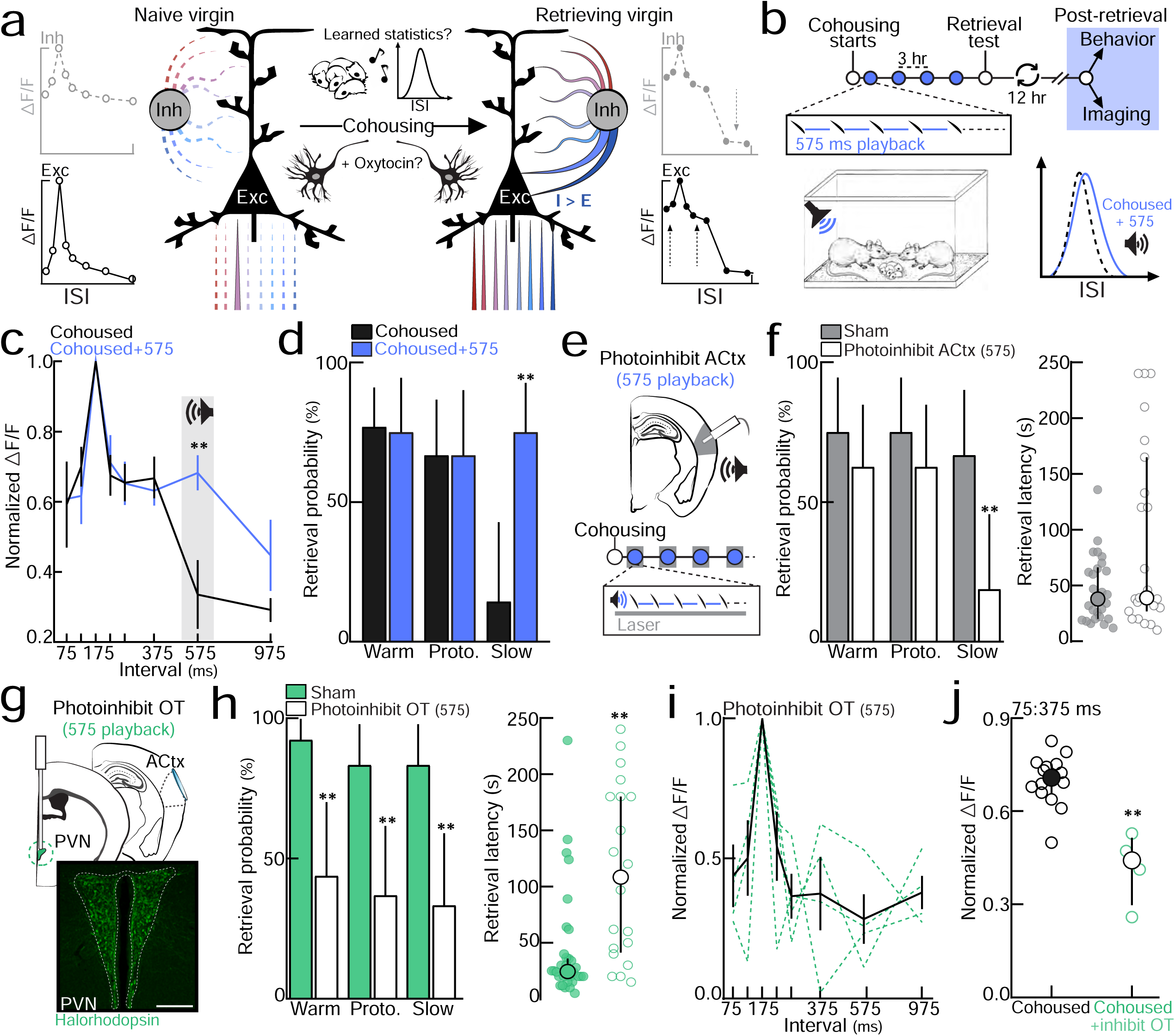
Auditory cortex and the oxytocin system are critical for learning about pup call features during cohousing. **a,** A schematic summarizing cortical changes during cohousing. **b,** Slow morphs (575 ms) were played during cohousing (‘cohoused+575’). Virgins were tested for pup retrieval or underwent two-photon imaging 24 hours after retrieval onset. **c,** Temporal tuning curves from virgins that heard 575 ms morphs during cohousing. Cohoused vs. cohoused+575: 575 ms, p=0.009. **d,** Virgins in the ‘cohoused+575’ group retrieved pups dubbed with slow morphs on significantly more trials than controls. Warm: cohoused (n=20/26 trials) vs. cohoused+575 (n=9/12 trials), p>0.99. Prototypes: n=12/18 vs. n=8/12, p>0.99. Slow: n=2/14 vs. n=12/16 trials, p<0.0001. Binomial test; Errors bars, 95% binomial CI. **e,** Photoinhibition of auditory cortex (ACtx) during 575 ms playback in cohoused virgins (‘photoinhibit-ACtx_(575)_’). **F**, Left-retrieval testing in sham and ‘photoinhibit ACtx_(575)_’ virgins. Warm: sham (n=9/12 trials) vs. photoinhibit-ACtx (n=10/16), p>0.99. Prototype: n=9/12 vs. n=10/16, p>0.99. Slow: n=8/12 vs. n=3/16, p<0.0001; Binomial test. Errors bars, 95% binomial CI. Right-retrieval latency on successful trials (p=0.20, Mann-Whitney test). Median±Interquartile. **g,** Halorhodopsin expression in oxytocin (OT) neurons in the PVN of the hypothalamus (scale bar, 250 µm). Playback and photoinhibition on same schedule as **b-f** (‘photoinhibit OT_(575)_’). **h,** Retrieval probability (left, p<0.0001. Binomial test; Errors bars, 95% binomial CI) and latency to retrieve (right, p=0.0003. Mann-Whitney test; Median±Interquartile) were reduced in the ‘photoinhibit OT_(575)_’ group. **i,** Temporal tuning in ‘photoinhibit OT_(575)_’ virgins. **j**, Temporal tuning across behaviorally-salient ISIs (p=0.001, Mann-Whitney test). Median±Interquartile. Cohoused = all experienced virgins imaged (N=14).

We then behaviorally tested these virgins for retrieval of cold pups dubbed over with slow morphs to determine whether they associated slower morphs with pup distress. Anesthetized pups were dubbed over with either prototypes (‘proto’) or 575 ms morphs (‘slow’) as described previously (**Fig. 1b**). Virgins that heard slow morphs during cohousing (‘Cohoused+575’) retrieved cold pups dubbed over with slow morphs (N=8 mice, n=12/16 trials) on significantly more trials than controls (N=9, n=2/14, p<0.0001; **Fig. 4d, Extended Data Fig. 12b**). Co-caring virgins can therefore learn to associate slower intervals with pup distress if heard frequently enough during initial maternal experience. Importantly, auditory cortical activity during the playback of slow morphs was required for virgins to learn this association (**Fig. 4e**, ‘photoinhibit-ACtx_(575)_’). Compared to sham animals (N=3 mice), optogenetic inhibition of auditory cortex during morph playback (N=4 mice) selectively prevented learning about slow ISIs (sham: n=8/12 trials, photoinhibit-ACtx_(575)_: n=3/16, p<0.0001; **Fig. 4f**-left). Overall, there was no difference in the latency to retrieve between the sham and optically-stimulated group (**Fig. 4f**-right). These experiments indicate that auditory cortex is a major site of plasticity specifically during exemplar learning. Not all sounds might be readily learned as distress calls though; virgins that heard single syllables (ISI:30 s) during cohousing were unable to associate these calls with isolated pups, suggesting an innate bias for a temporally-modulated bouts (**Extended Data Fig. 12c,d**).

Finally, we wondered what neural mechanisms underlie the re-tuning of virgin auditory cortex to represent relevant acoustic features for reliable pup retrieval. We focused on the peptide hormone oxytocin as this neurochemical is critical for maternal behavior^3, 8, 28^, and enables long-term plasticity in auditory cortex^8, 28, 29^. We hypothesized that perturbing oxytocin signaling exclusively during the playback of slow morphs (‘photoinhibit-OT_(575)_’) should also selectively abolish learning about slow ISIs. We cohoused naive virgins that expressed the inhibitory opsin halorhodopsin in oxytocin neurons (**Extended Data Fig. 13a**) with a dam and litter. We then optogenetically inhibited oxytocin neurons during the playback of 575 ms morphs on the same schedule as **Figure 4b-f**. Although photoinhibition only occurred during morph playback, auditory-based pup retrieval was perturbed overall compared to sham animals. The probability of retrieving vocalizing pups (sham (N=3 mice, n=12/13 trials) vs. photoinhibit-OT_(575)_ (N=5, n=7/16), p<0.0001), as well as cold pups dubbed over with prototypical calls (n=10/12 vs. 7/19, p<0.0001) or slow morphs (n=10/12 vs. 6/18, p<0.0001), was significantly reduced (**Fig. 4h-**left). Latency to retrieve was also significantly longer in the photoinhibition group (p=0.0003; **Fig. 4h**-ight).

Importantly, we observed that temporal tuning was also perturbed in the auditory cortex of virgins that received optical inhibition throughout cohousing (‘photoinhibit-OT_(575)_’, **Fig. 4i, Extended Data Fig. 13b**). Temporal tuning was significantly narrower across behaviorally-salient ISIs in optically-inhibited virgins (N=4 mice) compared to all the cohoused virgins imaged (N=14, p=0.001; **Fig. 4j**). This narrow tuning in experienced cortex was reminiscent of what we observed in experienced virgins that retrieved unreliably on 10-30% of trials (**Extended Data Fig. 6**), such that there was no significant difference in tuning width between these virgins (p=0.95; **Extended Data Fig. 13c**). These data are consistent with previous studies showing that oxytocin retunes cortical neurons and enhances spiking to acoustic stimuli^8, 29^. In line with our previous in vitro results^29^, we found that oxytocin significantly enhanced spike probability in response to a series of pulses in auditory cortical brain slices (**Extended Data Fig. 13d,e**). Oxytocinergic modulation may therefore act to transiently engage plastic mechanisms (e.g., via the reduction in intracortical inhibition), as well as on a long timescale such that chronic perturbations in the system will have long-lasting consequences on auditory cortical plasticity and maternal behavior.

Our results indicate that maternal behavior and auditory learning in mice results from interactions between an intrinsically primed cortex and experience-dependent learning processes. Previous studies showed that even during early postnatal development, rodent auditory cortex is especially sensitive to stimuli temporally-modulated around 5-6 Hz. This hardwired sensitivity could emerge from differential adaptation rates for call-evoked excitation and inhibition, and may serve as a mechanism by which pup-naive virgins are innately sensitized to prototypical calls for the rapid onset of maternal behavior. Auditory cortical activity in conjunction with oxytocinergic modulation, specifically in left auditory cortex, may then enable the restructuring of excitatory and inhibitory inputs to encompass feature distributions heard during cohousing. In this way, cortical neurons begin to generalize across variability in temporal statistics, enabling a robust response to potentially any vocalization within the pup call category for reliable parental care in noisy or complex environments.

## Methods

### Stimulus library and vocalization analysis

Pup vocalizations were recorded from isolated pups (postnatal day (PND) 1-7) using an ultrasonic microphone (Avisoft Bioacoustics CM16/CMPA, sampling rate: 200 kHz). Analysis of ISIs was performed in Adobe Audition. 4,540 inter-syllable intervals were measured from 32 audio recordings (2:50-3:30 min. each, total time: ∼96 min.) and a frequency distribution was generated (**Fig. 1a**). Pup calls used in imaging and behavioral experiments were de-noised and matched in peak amplitude (Adobe Audition). Prototypical calls contained 4-5 syllables and had an average ISI of 150-200 ms (bin:175±25 ms). Prototypes were morphed in the temporal domain by either adding (+50, +100, +200, +400, +800 ms) or subtracting (–50, –100 ms) time between syllables to generate a morph set for each prototypical call. This resulted in bins sized ±25 ms around the bin center: 75, 125, 225, 275, 375, 575, and 975 ms (**Extended Data Fig. 1**).

### Behavior and cohousing

6-48-week-old C57BL/6 virgin females (Taconic Biosciences and Jackson Laboratory) were used in all experiments. Lactating dams (∼3-5 months old) with pups ranging from postnatal days 1-7 were used for cohousing and behavioral testing. Prior to experimentation, all virgins were tested for retrieval to establish a baseline as described previously^8^. Briefly, females were given 15-30 minutes to acclimate to a novel behavioral arena (38×30×15 cm). At least three pups were placed in the corner surrounded by nesting material. A trial was initiated by placing one pup in the far corner. Females were given two minutes to retrieve the pup back to the nest. If virgins failed to retrieve, the pup was placed back in the nest and another trial was initiated for a total of 5-10 trials. Virgins that did not retrieve on any trials were housed with age-matched cagemates (‘naive virgin’) until further testing. Experienced virgins were housed with a dam and litter for at least 72 hours before further experimentation.

To test retrieval of anesthetized pups dubbed over with auditory stimuli (**Fig. 1b,c, Extended Data Fig. 2, Fig. 4d-h, Extended Data Fig. 12c**), experienced females were housed with pups in a behavioral arena (38×30×15 cm), modified with an electrostatic speaker in the far corner and an adjacent door. Animals were given 12-24 hours to acclimate to the arena. When testing a cohoused virgin, the dam was removed prior to testing and vice versa. All testing was done under red-light conditions. To begin, 3-5 pups were removed from the nest and anesthetized on ice for 10-15 minutes to prevent vocalizations. A trial was initiated when the female remained in the nest for at least two minutes. The corner door was then opened and a warm or cold/anesthetized pup was placed in front of the adjacent speaker. On warm trials, pups vocalized as expected (verified by ultrasonic microphone (Avisoft Bioacoustics)). On cold pup trials, no stimuli were played and an ultrasonic microphone was used to ensure no vocalizations were emitted from anesthetized pups. On trials in which cold pups were dubbed over with stimuli, a series of 5-8 prototypical calls or morphs were emitted from the corner speaker (inter-bout interval: 2-3 s). For single syllable trials, one syllable taken from prototypical calls was played every 30 seconds. Mice were given four minutes to retrieve the warm or cold pup back to the nest. If the pup was retrieved, the latency to retrieve was recorded. If the pup was not retrieved, the trial was marked a failure and another trial was initiated upon the mouse remaining in the nest for at least two minutes. Testing continued until each animal performed 8-16 trials, or testing could not continue as the result of nursing, cannibalization, or failure to enter the nest after three hours. Trials in which cold pups began vocalizing or warm pups failed to vocalize were excluded from analysis. Data from **Fig. 1c** and **Extended Data Fig. 2** were collected from lactating dams and experienced virgins to establish which ISIs elicited retrieval due to experimental limitations. Both dams and experienced virgins had similar behavioral responses (**Extended Data Fig. 2a**). All behavioral testing following long-term cohousing manipulations (i.e., pup call exposure in **Fig. 4, Extended Data Fig. 12**) was exclusively performed in virgins.

In order to assess behavioral preferences for pup calls (**Extended Data Fig. 3**), dams and their litters, along with cohoused virgins, were placed in a modified y-maze box (Plexiglas; 47×36×28 cm) 24 hours prior to testing. When testing a cohoused virgin, the dam was removed prior to testing and vice versa. At the start of each trial, three pups were placed in the left, right, and center chambers. After retrieval of all three pups, the speakers in the left and right chambers were simultaneously switched on to play competing pup call bouts (inter-bout interval: 1 second). A prototypical call was played from one speaker, while the same call morphed in the temporal domain was played from the other. The calls played continuously until a chamber was chosen or the timeout period (two minutes) elapsed. Prototypical and morphed calls alternated left and right speakers across trials. Room entry (% trials) was calculated by dividing the number of approaches by the total number of trials a given ISI was played.

In the operant conditioning paradigm (**Fig. 1i-k, Extended Data Fig. 8**), pup-naive virgins could hold down a lever to turn off continuously playing pup call sounds. The operant chamber (25×20×15 cm) consisted of an electrostatic speaker and metal lever connected to a capacitive touch sensor (SparkFun). Pup call presentation was controlled by custom-written programs in MATLAB interfacing with an RZ6 Multi-I/O processor (Tucker-Davis Technologies). Naive virgins were in one of three groups, listening to either prototypical pup calls (ISI:150-200 ms), fast morphs (ISI:50-100 ms), or slow morphs (ISI:550-600 ms). At the start of a session, virgins were placed in the arena and immediately calls or morphs began continuously playing. Stimulus playback was only paused via lever press. Brief touches resulted in a pause in stimulus presentation for a set duration, which scaled with the ISI duration to control for discrimination between the inter-syllable interval and pause across groups (15-20x ISI duration; prototypes: ∼3s, fast: ∼1.5s, slow: ∼9s). For example, in the prototype group a brief lever press for less than 3 seconds resulted in a pause of prototype playback for 3 seconds. As press duration increased beyond this threshold, playback was paused for the entire duration of the press. Only presses ≥1 ms were counted as touches. Learning trajectories were smoothed using a moving average (bin size: 2 days) for plotting purposes in **Fig. 1j.**

### Cranial window implantation & head-posting

All procedures were approved under NYU Langone Institutional Animal Care and Use committee protocols. For two-photon calcium imaging, cranial window implantation over left or right auditory cortex was performed as previously described^20^. Females were anesthetized with isoflurane (1.0-2.5%) and a 3 mm craniotomy was centered 1.75 mm anterior to the lambda suture. Dexamethasone (0.01-0.025 mL) was injected subcutaneously to reduce intra-cranial swelling. Adeno-associated viruses encoding GCaMP6f (0.75-1.0 μL) were injected in the center of the craniotomy at a depth of ∼1000 μm using a 5 μL syringe (33 gauge needle, Hamilton). To restrict expression of GCaMP6f to excitatory neurons, we used AAV9.CamKII.GCaMP6f (UPenn Vector Core or Addgene). For expression of GCaMP6f in inhibitory interneurons, we injected rAAV.mDLX.GCaMP6f^24^ (gift of Jordane Dimidschstein and Gordon Fishell) into wild type mice or injected AAV1.Syn.Flex.GCaMP6f (UPenn Vector Core or Addgene) into Gad2-Ires-Cre C57BL/6J mice (Jackson Laboratory). A 3 mm glass coverslip was secured over the craniotomy using a mixture of Krazy Glue and Acrylic resin powder (Lang Dental Manufacturing). For head fixation during in vivo calcium imaging and in vivo whole-cell recordings, custom-made headposts (Ponoko) were then secured to the skull with C&B Metabond dental cement (Parkell). Animals were given 2-4 weeks for viral expression and recovery.

### In vivo two-photon calcium imaging

Two photon-calcium imaging was performed in awake, head-fixed mice as previously described^20^. Animals were habituated to head-fixation 15-30 minutes a day for 2-3 days prior to data acquisition. We used a multiphoton imaging system (Sutter Instruments) and a 900 nm Ti:Sapphire laser (MaiTai, Spectra-Physics) to obtain GCaMP6f signals in auditory cortex. Images from layer 2/3 regions (∼300 μm^2^, 256×256 pixels) were collected using ScanImage (HHMI, Vidrio Technologies) at a rate of 4 Hz (0.26 s/frames). This scanning rate did not affect our ability to detect responses to stimuli faster than 4 Hz (**Extended Data Fig. 4d**). During image acquisition, auditory stimuli were played through an electrostatic speaker (10-12 cm from the contralateral ear) connected to an RZ6 Multi-I/O processor and controlled by RPvdsEx (Tucker-Davis Technologies) interacting with ScanImage (HHMI). Half octave pure tones ranging 4-64 kHz (500 ms, 10 ms cosine ramp) at 70 dB sound pressure level (SPL) were played in a pseudorandom order (repetition rate: 0.2 Hz). For temporally-modulated pure tones, a tone was selected based on the best frequency for the region (**Extended Data Fig. 7**) and a series of pure tone pips at this frequency (70 dB SPL, 50-80 ms, 2 ms cosine ramp) were then played with the following ISIs: 75, 175, 375, and 575 ms (binned±25 ms).

For pup call presentation, a library of 6-9 prototypical calls (2 ms cosine ramp, ISI:175±25 ms) were played in a pseudorandom order (repetition rate: 0.1 Hz). Acquired signals were screened offline to determine the presence of call-responsive neurons, and which calls evoked responses in those neurons. Temporally morphed stimuli were then played exclusively for those prototypical calls. For chronic two-photon imaging experiments (**Fig. 3**), cohoused virgins were tested for retrieval every 12 hours. Every other time point, retrieval testing was followed immediately by calcium imaging. Neuronal responses were always assessed at first retrieval and 24 hours after first retrieval. Retrieval onset ranged from 12–109 hours of cohousing in n=13 virgins. To control for exposure to calls every day under the two-photon, pup-call tuning was measured every 24 hours for three consecutive days in non-cohoused virgins (**Extended Data Fig. 11c,d**).

### Image processing and analysis

Images were aligned frame-by-frame to a stable set of frames using TurboReg for x-y movement correction (Fiji/Image J, NIH). Regions of interest (ROIs) were manually drawn on an average image for extraction of raw calcium signals (Fiji/Image J, NIH)^20^. Cells were deemed as responsive to any of the 6-9 prototypes if the integral of the fluorescence signal during the stimulus epoch (starting 250 ms post-call onset, 1 s in duration) significantly increased from baseline (‘F_0_’, 1 s prior to stimulus onset; p<0.05, Student’s paired one-tailed t-test). Neurons responsive to prototypical calls were then assessed for their responses to temporally-morphed calls. ΔF/F (%) was calculated as the average change in fluorescence during the stimulus epoch: 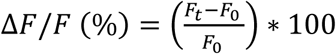. For temporal morphs, frames were added or subtracted to the stimulus epoch for each 500 ms added or subtracted to the total call duration, respectively. This window was shifted according to the peak of the signal for all stimuli on a cell-by-cell basis.

To measure the similarity in evoked responses between a prototype and its morph, normalized ΔF/F was calculated as: 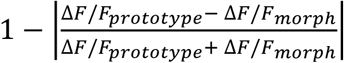. The slope of the tuning curve at the transition between behaviorally-salient (75:375 ms) and non-salient (575:975 ms) ISIs was calculated by taking the slope of the line between 375:975 ms. In chronic two-photon imaging experiments, spatial correlations were used to assess whether or not a prototype-responsive cell was present over several days, as previously described^27^. Briefly, following manual inspection, a 100×100-pixel image surrounding a given ROI was taken for each imaging session. Spatial correlations were calculated for a given ROI across days (‘within ROIs’) and with other ROIs in the population (‘across ROIs’). Cells with spatial correlations below the 95^th^ percentile for the null distribution (‘across ROIs’) were eliminated from the correlation analysis in **Fig. 3b-e**.

To assess whether neuropil contamination had any significant effect on call-evoked ΔF/F values, we performed corrections on two data sets as described previously^26^. Briefly, the true cell body fluorescence was calculated as: *F_true_*(t) = *F_raw_*(t) − r * *F_neuropil_*(t), in which (t) is time, *F_raw_* is the raw fluorescence averaged within an ROI, *F_neuropil_* is the neuropil fluorescence surrounding a given ROI, and r is a ratio of contamination 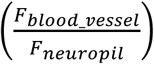. Neuropil correction had no significant effect on call-evoked ΔF/F values in excitatory or inhibitory neurons (**Extended Data Fig. 4e-g**).

Finally, we calculated ΔF/F for pure tone-responsive neurons similarly to our pup calls. For temporally-modulated pure tones (**Extended Data Fig. 7**), only cells that had statistically significant responses to the best frequency and the 5 Hz tone sequence (the prototypical repetition rate) were included in further analysis (p<0.05, Student’s paired one-tailed t-test). Normalized ΔF/F was calculated similarly to pup calls, such that the response evoked by each temporally-modulated sequence was normalized to the response evoked by the 5 Hz temporally-modulated tones (ISI: 175 ms).

### In vitro whole-cell recordings

In vitro recordings were performed in acute slices of auditory cortex prepared from pup-naive C57Bl/6 wild-type mice. Animals were deeply anesthetized with 5% isoflurane and decapitated. The brain was rapidly placed in ice-cold dissection buffer containing (in mM): 87 NaCl, 75 sucrose, 2.5 KCl, 1.25 NaH_2_PO_4_, 0.5 CaCl_2_, 7 MgCl_2_, 25 NaHCO_3_, 1.3 ascorbic acid, and 10 D-Glucose, bubbled with 95%/5% O_2_/CO_2_ (pH 7.4). Slices (250–300 µm thick) were prepared with a vibratome (Leica P-1000), placed in warm artificial cerebrospinal fluid (ACSF, in mM: 124 NaCl, 2.5 KCl, 1.5 MgSO_4_, 1.25 NaH_2_PO_4_, 2.5 CaCl_2_, and 26 NaHCO_3_,) (33-35°C) for <30 min, then cooled to room temperature (22-24°C) for at least 30 minutes before use. Slices were then transferred to the recording chamber and superfused (2.5-3 ml/min) with oxygenated ACSF at 33°C. Somatic whole-cell voltage-clamp or current-clamp recordings were made from layer 2-3 pyramidal cells with an Multiclamp 200B amplifier (Molecular Devices) using IR video microscopy (Olympus). Data were filtered at 2 kHz, digitized at 10 kHz, and acquired with Clampex 10.7 (Molecular Devices). Data were analyzed using custom-written Matlab code (MathWorks) and Clampfit 10.7 (Molecular Devices).

In order to assess short-term plasticity of EPSCs and IPSCs, voltage-clamp recordings were acquired while focal extracellular stimulation (0.5 ms, 3-700 μA) was applied with a bipolar glass electrode at a rate of 1 stimulation every 75, 175, or 575 ms (bin±25 ms). Patch pipettes (3-8 MΩ) were filled with the following intracellular solution: 130 mM Cs-methanesulfonate, 1 mM QX-314, 4 mM TEA-Cl, 0.5 mM BAPTA, 4 mM MgATP, 0.3 mM Na-GTP, 10 mM phosphocreatine, 10 mM HEPES, pH 7.2). EPSCs were acquired at −70 mv, while IPSCs were acquired from −40 to 0 mV. The peak amplitude of evoked EPSCs and IPSCs were measured and normalized to the event evoked by the first stimulation on each trial (S_n_ / S_1_; **Fig. 1h**).

To assess the modulatory effects of oxytocin (**Extended Data Fig. 13e,e**), whole-cell recordings were acquired in current-clamp configuration with patch pipettes (4-8 MΩ) containing the following intracellular solution (in mM): 130 Cs-methanesulfonate, 4 TEA-Cl, 10 Phosphocreatine, 0.5 EGTA, 10 HEPES, 1 QX-314, 4 MgATP, 0.3 NaGTP, pH 7.2. Cells were injected with current to raise the membrane potential near spiking threshold while focal extracellular stimulation was applied with a bipolar glass electrode at a rate of 1 stimulation every 550-575 ms. Threshold was empirically determined for each cell as the membrane potential that resulted in evoked action potentials ≤ 40% of trials. Once threshold was determined, a stable baseline was established for 5-10 minutes. 1 *μ*M oxytocin (Tocris) in ACSF was washed on for 10-15 minutes, followed by a washout period. To control for continuous extracellular stimulation, the same protocol was run in the absence of oxytocin. Spike probability was calculated as the probability of evoking a spike in response to a given stimulation 15-30 minutes following wash onset or post-baseline acquisition in the case of controls.

### In vivo whole-cell recordings

Animals were anesthetized with 1.5-2.0% isoflurane and head-fixed using custom-made stainless steel headbars (Ponoko). A small craniotomy was performed over left auditory cortex and whole-cell voltage-clamp recordings were obtained from layer 2/3 (200-400 µm from the pial surface) with a Multiclamp 700B amplifier (Molecular Devices). Borosilicate glass pipettes (Sutter) with resistance 5-7 MΩ contained (in mM): 130 Cs-methanesulfonate, 1 QX-314, 4 TEA-Cl, 0.5 EGTA, 4 MgATP, 0.3 NaGTP, 10 phosphocreatine, 10 HEPES, pH 7.2. Ri: 206.20±57.09 MΩ (s.d.). Once whole-cell configuration was obtained, five prototypical calls were played to determine the best prototypical call for each cell (i.e., the call that evoked the maximal EPSC). Once the best call was determined, the prototype was played in conjunction with a set of temporal morphs (125, 225, 575 ms). Cells were held at −70 mV for excitatory currents, and between 0 to +40 mV for inhibition. Recordings were analyzed using Clampfit. Only neurons that exhibited at least five significant prototype-evoked E/IPSCs were used for further analysis. Synaptic responses were calculated as the PSC area (mv*ms) divided by the duration of the predetermined analysis window (ms); the baseline analysis window was 1 second, and prototypical and morphed calls were analyzed from the start of the call to 250 ms following call offset. Absolute magnitudes (pA) from the largest 5-10 evoked trials were used for analysis. Evoked synaptic responses were calculated by subtracting the baseline response from the stimulus response (*PSC_stimulus_* − *PSC_baseline_*). The relative difference in the evoked IPSC and EPSC (‘I-E ratio’) within a neuron was calculated for each stimulus as 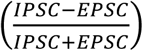.

### Pup call exposure

Non-retrieving virgins were cohoused with a dam and litter. An electrostatic speaker was placed ∼15 cm above the nest. 30-60 minutes after the beginning of cohousing, auditory stimuli were played every ∼3-4 hours for 12-hours, interleaved with 12-hour blocks without stimuli. This was meant to mimic long blocks of silence followed by brief periods of vocalizing observed during long-term audio recordings. Virgins were tested for retrieval every 12 hours. For 575 ms exposure (**Fig. 4b-j, Extended Data Fig. 13b,c**), a set of 5-8 morphed calls (ISI:550-600 ms, ‘575 ms’) were repeated 10-20 times each in a pseudorandom order (one morph presented every 8-12 seconds). For single syllable exposure (**Extended Data Fig. 12c,d**), a series of single syllables was presented with 30 seconds between each syllable. All virgins underwent two-photon calcium imaging and/or behavioral testing in the cold pup retrieval test (described above, Methods: Behavior and cohousing) 24 hours after retrieval onset.

### Inhibitory optogenetics

Surgical preparation was performed as described above (Methods: Cranial window implantation). For optogenetic inhibition of auditory cortex (**Fig. 4e,f**), opsins (0.75-1.0 μL) were injected in the center of a small craniotomy (1.75 mm anterior to the lambda suture) at a depth of ∼1000 μm using a 5 μL syringe (33-gauge needle, Hamilton). The auditory cortex was inhibited in two ways: 1) the hyperpolarizing opsin Halorhodopsin (eNpHR3.0) was expressed in excitatory neurons of wildtype C57BL/6J virgins (AAV1.CaMKIIa.eNpHR.EYFP; Addgene), or 2) the depolarizing opsin Channelrhodopsin (AAV1.EF1a.DIO.hChR2.EYF; Addgene) was expressed in a cre-dependent manner in interneurons using Gad2-IRES-Cre C57BL/6J mice (Jackson Laboratory). A 2 mm long optic fiber (Ø400 µm Core, Ø1.25 x 6.4 mm Ceramic Ferrule, 0.39 NA; ThorLabs) was then implanted at a depth of 250-400 μM and secured to the skull with C&B Metabond dental cement (Parkell). Animals were given 2-4 weeks for viral expression and recovery, followed by habituation to the patch cable (Ø400 µm Core, 0.39 NA FC/PC to Ø1.25 mm; ThorLabs) for 3-5 days prior to beginning cohousing. Implanted virgins were cohoused and exposed to slow morphs (ISI:575 ms) on the same schedule as described in **Fig. 4b**. During these exposure timepoints, the opsins were stimulated to inhibit the auditory cortex during call playback (Halorhodopsin: 532 nm wavelength, 1–3 mW/mm^2^; Channelrhodopsin: 473 nm wavelength, 1-3 mW/mm^2^). Sham animals included virgins injected with halorhodopsin (AAV1.CaMKIIa.eNpHR.EYFP; Addgene; N=1) that received no optical stimulation, or virgins injected with the structural marker YFP (AAV.CamKII(1.3).eYFP; Addgene; N=2) that received optical stimulation (532 nm wavelength, 1–3 mW/mm^2^). Behavioral testing in the cold pup retrieval assay was performed 24 hours following retrieval onset.

Optogenetic inhibition of oxytocin neurons was performed similarly to cortical inhibition (**Fig. 4g-j, Extended Data Fig. 13a-c**). Briefly, 1.0-1.5 μL of a cre-dependent halorhodopsin (AAV5.Ef1a.DIO.eNpHR.EYFP; Addgene) was injected via Hamilton syringe in to the left paraventricular nucleus of hypothalamus (AP: −720µM, ML: +120 µM; DV: 4,500-4,750 µM) of Oxt-Ires-Cre C57BL/6J mice (Jackson Laboratory). Transgenic mice expressing halorhodopsin in oxytocin^+^ neurons were also used (Oxt-Ires-cre x Ai39 mice; Jackson Laboratory). In a subset of mice, GCaMP6f (AAV1.Syn.Flex.GCaMP6f; Addgene) was also injected in auditory cortex as described above (Methods: Cranial window implantation). Sham animals were injected with a cre-dependent YFP as a control (AAV5.Ef1a.DIO.EYFP; Addgene). A 5 mm optic fiber (Ø200 µm Core, Ø1.25 x 10.5 mm Ceramic Ferrule, 0.39 NA; ThorLabs) was then implanted at a depth of ∼4,250-4,500 µM and secured to the skull with C&B Metabond (Parkell). During cohousing, oxytocin neurons were inhibited during call playback on the same schedule as in the cortical inhibition experiments (532 nm wavelength, 1–5 mW/mm^2^). Behavioral testing and/or two-photon imaging were performed 24 hours following retrieval onset.

Viral expression was confirmed using immunohistochemistry (**Fig. 4g**). Briefly, animals were perfused with 4% paraformaldehyde following experiments. Brains were removed and post-fixed in 4% PFA for 24 hours at 4°C, followed by immersion in 30% sucrose for 48 hours at 4°C. Brain were embedded in Optimal Cutting Temperature compound and stored at −80°C prior to sectioning. 50 µm thick slices were cut using a cryostat and stained using standard immunohistochemistry histological methods. Primary antibody (1:500): Rabbit anti-GFP (AB290, Abcam). Secondary antibody (1:1000): Goat anti-rabbit, Alexa Fluor® 488 (AB150077, Abcam).

### Statistics

Power analysis was performed for behavioral experiments to determine sample size for statistical significance with a power of 0.8. Y-maze experiments required at least n=20 trials (**Extended Data Fig. 3**) and the cold pup retrieval assay required at least n=7 trials (**Fig. 1c, Extended Data Fig. 2, Fig. 4d,f,h, Extended Data Fig. 12c,d**). These requirements were satisfied in terms of the total number of trials for each experiment. For whole cell recordings *a priori* power analysis was not performed, however post-hoc analysis was used to ensure we obtained a power ≥ 0.8 for experiments in **Fig. 2g-m**. For sample size reporting, the number of mice is reported for across animal comparisons. For comparisons across neurons, the total number of neuronal tuning curves is reported as a given neuron may have responded to multiple calls. Sample size is only reported the first time a group is mentioned within a figure legend. Data was repeated in the manuscript for appropriate comparisons as follows: 1) Experienced virgins from **Fig. 1c** were used in **Fig. 4d** and **Extended Data Fig. 12c,d** as ‘Cohoused’ virgins, 2) virgins from **Fig. 2h** (24 hours post-retrieval) were used in **Fig. 4c** (Note: **Fig. 2h** depicts single-cell tuning, **Fig. 4c** depicts each virgins’ tuning curve). One or two-tailed student t-tests were performed appropriately based on hypothesis testing. Unless otherwise stated, a student’s unpaired two-tailed t-test was used. The Bonferroni method was used to correct for multiple comparisons when appropriate for all ANOVA testing. Error bars and shading on line plots denote ±s.e.m unless otherwise stated. * represents p<0.05, ** represents p<0.001 throughout the manuscript.

---

**Extended Data Figure 1.**
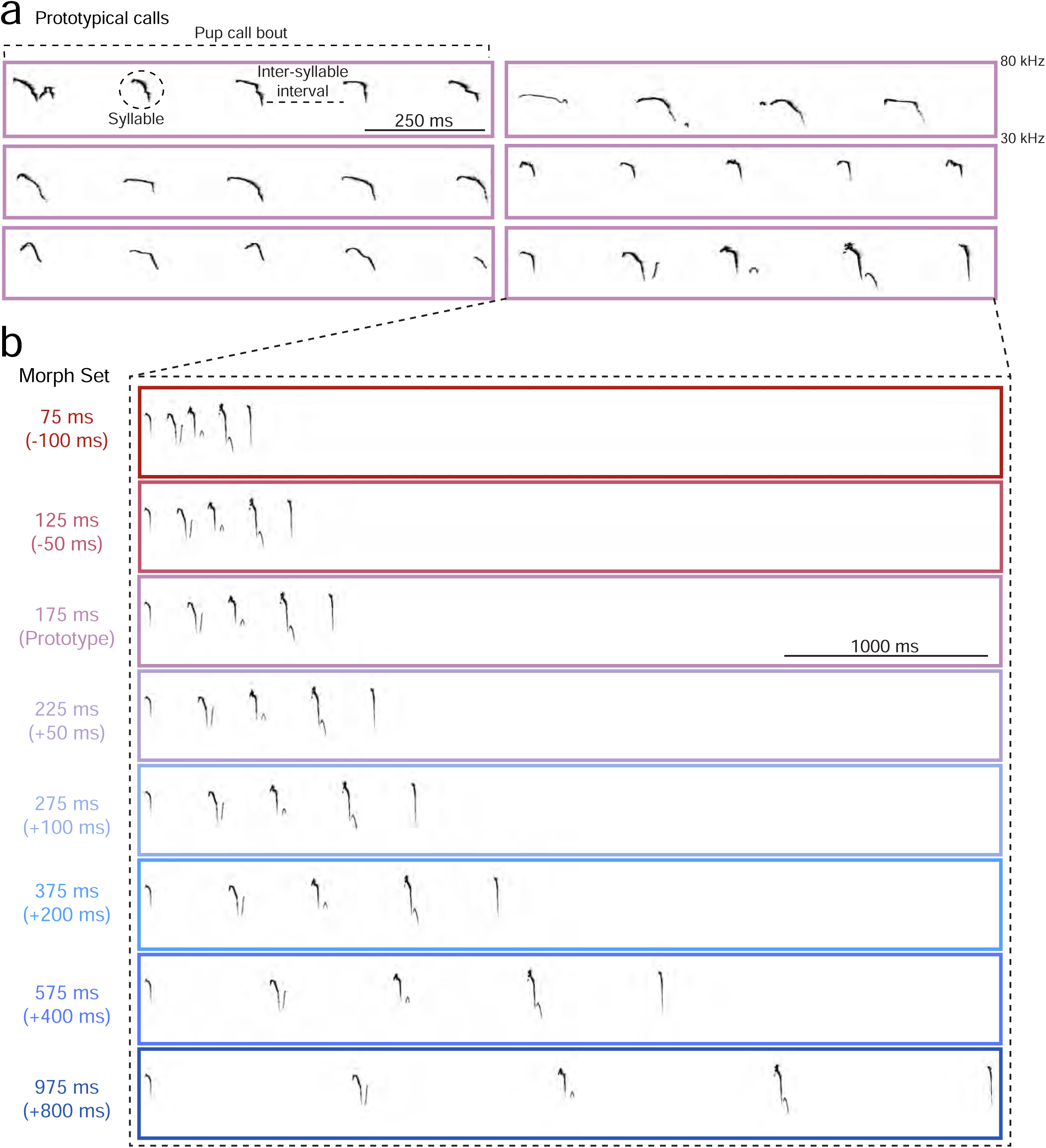
Stimulus library of prototypical and morphed pup calls. **a,** A set of six prototypical pup calls were selected from a library of pre-recorded calls. Prototypical calls had an average ISI of 150-200 ms (Bin:175±25 ms). **b,** Example of one prototypical call morphed in the temporal domain. Time was added or subtracted from the ISI to slow or speed up the calls, respectively. Other features (e.g., frequency content) remained the same across all temporal morphs. A set of seven morphs was generated for each prototypical call (Bin center: 75, 125, 225, 275, 375, 575, 975 ms), resulting in a library of 42 pup call sounds. Bin size: ±25 ms. Color indicate overall speed/duration of ISIs, with red representing calls with shorter ISIs and blue representing slower calls with longer ISIs.

**Extended Data Figure 2.**
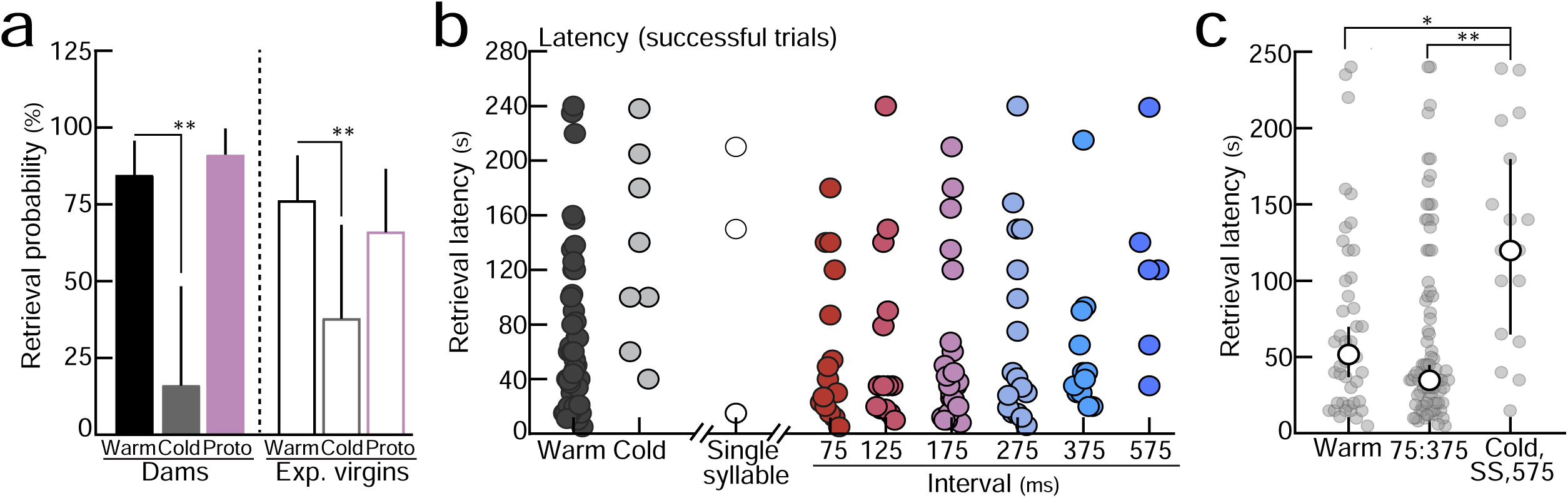
Retrieval of anesthetized pups dubbed over with prototypes and temporal morphs. **a,** Breakdown of retrieval behavior in the cold pup retrieval assay across dams and experienced virgins from Fig. 1c. Both groups exhibited similar behavioral response profiles. Dams (N=5-16 mice) and experienced virgins (N=9-17) retrieved warm pups and cold pups dubbed over with prototypical calls, but not cold (anesthetized) pups. Dams: warm (n=23/27 trials) vs. cold (n=3/12), p<0.0001, vs. prototype (n=11/12), p=0.97. Experienced virgins: warm (n=20/26) vs. cold (n=5/13), p=0.003, vs. prototype (n=12/18), p=0.26 (Binomial test). Error bars, 95% binomial CI. **b,** Latency to retrieve on successful retrieval trials from Fig. 1c. **c,** Latencies from **Extended Data Fig. 2b** binned based on retrieval probability in Fig. 1c. On successful retrieval trials, the latency to retrieve cold pups and pups dubbed over with 575 ms or single syllable (‘SS’) morphs was significantly longer than warm pups (p=0.02) and pups dubbed over with morphs that reliably elicited retrieval (75:375 ms, p=0.001; Kruskal–Wallis H-test). Error bars, median±95% CI.

**Extended Data Figure 3.**
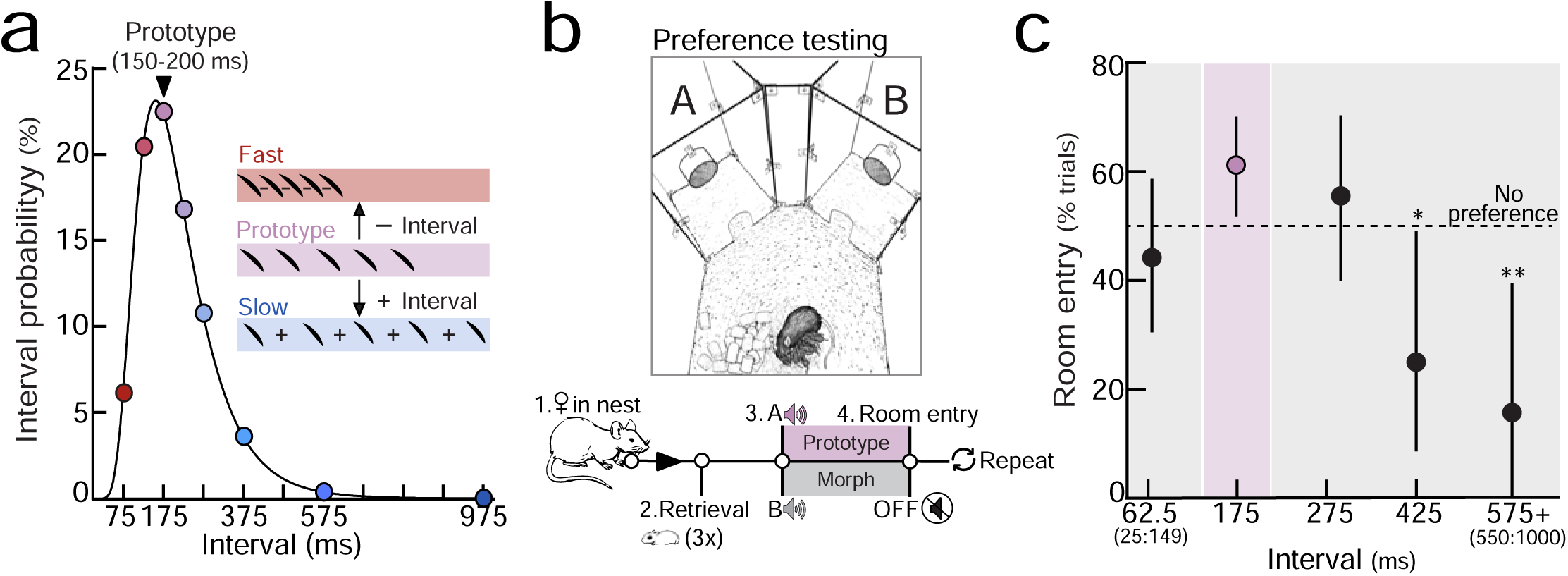
Experienced females approach speakers playing pup calls and temporal morphs. **a,** Same distribution as Fig. 1a**. b,** A y-maze was used to assess approach towards speakers playing pup calls and morphs in the absence of a live pup. Speakers in one room played a prototypical pup call (ISI:175 ms), while a competing speaker in the other room played a spectrally-matched, temporal morph. Mice were given two minutes to enter a room. **c,** When competing morphs contained ISIs 25:149 ms (62.5 ms, n=23/52 trials) or 201:350 ms (275 ms, n=25/45), experienced females showed no significant preference between the prototype and morphs (62.5 ms, p=0.50; 275 ms, p=0.46). Mice showed a significant preference for prototypes when ISIs were slower than 350 ms (425 ms (n=5/20), p=0.04; 575+ ms (n=3/19), p=0.005). Binomial test; Error bars, 95% binomial CI. Data binned ±75 ms (except 575+).

**Extended Data Figure 4.**
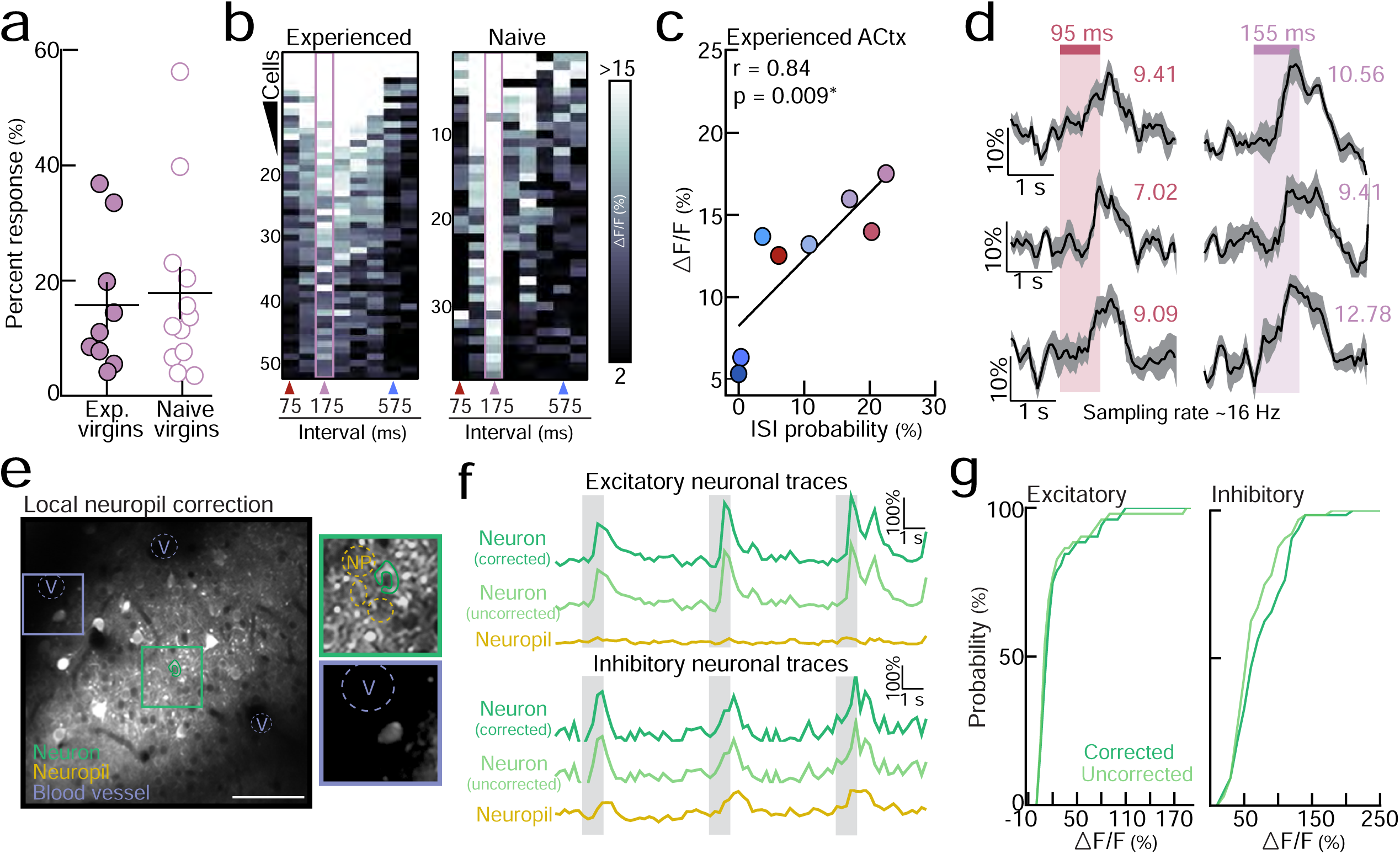
Two-photon calcium imaging of auditory cortical responses to pup calls. **a,** Percentage of prototype-responsive excitatory neurons in experienced (N=9 mice) and naive virgins (N=12; p=0.74). **b,** Sample heat maps of all the prototype-responsive neurons in an imaging region from an experienced (left) and naive (right) virgin. Color gradient denotes ΔF/F (%). Note strong preference for prototypes in naive cortex (pink bar). **c,** In experienced virgins, ΔF/Fs were correlated with ISI probability from distribution in Fig. 1a (Pearson’s r=0.84, p=0.01). Colors reflect ISI bins reported in Fig. 1a and **Extended Data** Fig. 1. **d,** Example ΔF/F Ca^2+^ traces acquired at 16 Hz from an experienced virgin. ΔF/F (%) is indicated by the colored number above each trace. All three prototype-responsive neurons have similar evoked responses between the fast morph (ISI:95 ms) and prototype (ISI:155 ms), consistent with neuronal responses acquired at 4 Hz (see Fig. 1e). **e-g,** Neuropil correction on sample data sets (N=1 imaging region for excitatory neurons; N=1 imaging region for inhibitory neurons). **e,** Fluorescence was corrected by measuring background fluorescence in vasculature (‘V’, purple) and neuropil surrounding each ROI (‘NP’, yellow). See methods. **f,** Example ΔF/F Ca^2+^ traces from an excitatory neuron (top) and an inhibitory neuron (bottom). **g,** Neuropil correction on sample data had no significant effect on prototype-evoked ΔF/F (%) in excitatory neurons (n=52 neurons, p=0.60) or inhibitory neurons (n=64 neurons, p=0.14; Kolmogorov-Smirnov test).

**Extended Data Figure 5.**
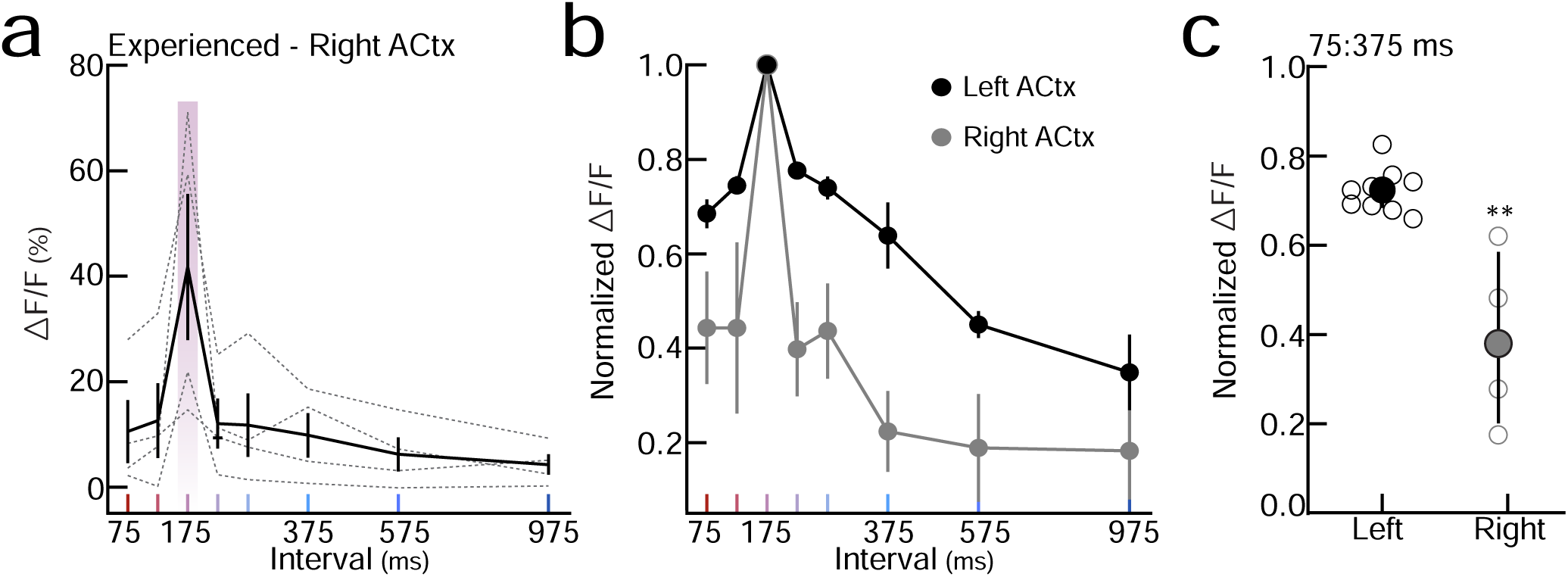
Left, but not right, auditory cortex is broadly tuned across temporal statistics in experienced females. **a,** Excitatory neuronal tuning in right auditory cortex of experienced virgins (N=4 mice). **b,** Excitatory neuronal tuning normalized to prototypes in the left (black) and right (gray) auditory cortex of experienced virgins (N=9 mice, from Fig. 1g). **c,** Response similarity between the prototype and morphs (‘Normalized ΔF/F’) was significantly reduced across behaviorally-salient ISIs (75:375 ms) in right auditory cortex (p=0.003; Mann-Whitney test). Error bars, median±interquartile range.

**Extended Data Figure 6.**
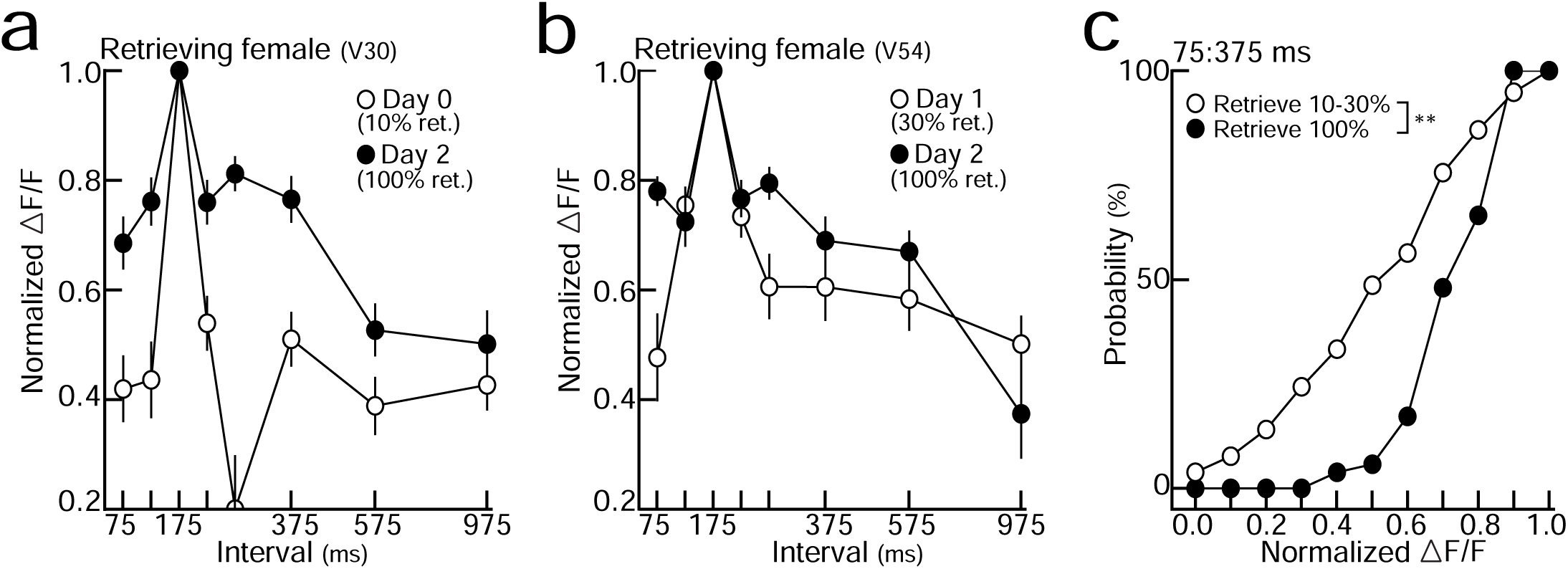
Reliable retrieval is associated with broad temporal tuning in experienced females. **a,b,** Two example experienced virgins that exhibited unreliable retrieval behavior on 10% (left) and 30% (right) of trials in the standard pup retrieval test (see methods). Temporal tuning at baseline (open circles) broadened following the onset of reliable retrieval behavior (100% of trials, closed circles). ‘Days’ denote days of cohousing with a dam and litter. **c,** Cumulative distribution of normalized ΔF/Fs averaged across behaviorally-salient ISIs (75:375 ms) before (N=2 mice, n=78 neurons) and after the onset of reliable retrieval (N=2, n=52). A larger proportion of neurons were more broadly tuned (higher normalized ΔF/Fs) when experienced virgins retrieved on 100% of trials (p<0.0001, Kolmogorov-Smirnov test).

**Extended Data Figure 7.**
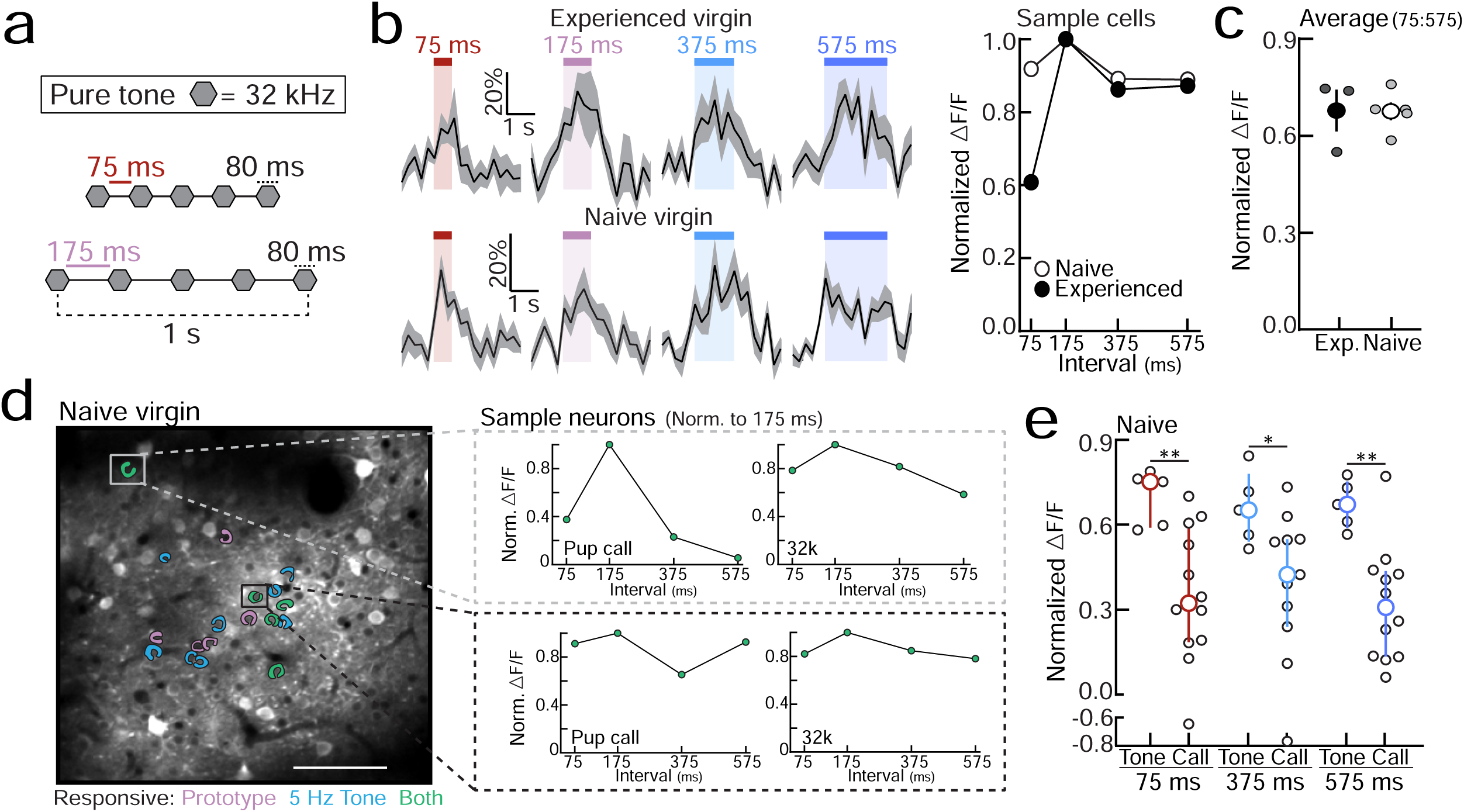
Auditory cortical neurons are broadly tuned across temporally-modulated pure tones in naive cortex. **a,** The best frequency for a given imaging region was selected and used to assess cortical tuning across temporally-modulated pure tone sequences. For example, a stimulus set consisted of five tone pips (32 kHz, 80 ms) separated by the following ISIs: 75, 175, 375, or 575 ms. **b,** Left, example ΔF/F Ca^2+^ traces evoked by temporally-modulated sequences of 32 kHz tones. Note sample cell from naive cortex, which responds robustly across all ISIs. Right, sample cell quantification. ΔF/F (%) normalized to the response evoked by the prototypical repetition rate (ISI:175 ms). **c,** ΔF/Fs evoked by temporally-modulated tones were normalized to the 175 ms ISI stimulus and averaged across all stimuli as a measure of tuning width. No significant difference between experienced (N=3 mice, n=50-94 neurons) and naive (N=4, n-31-45) females (p=0.97). **d,** Sample imaging region from a naive virgin. Prototypical pup calls (pink) and temporally-modulated 32 kHz tones with ISIs ∼175 ms (blue, ‘5 Hz tone’) activated a small subset of the same cells (green). Neurons responsive to both stimuli could have distinct tuning across ISIs (inset). **e,** Normalized ΔF/F values to temporally-modulated tones (N=5 mice) and pup calls (n=11-12) in naive auditory cortex. Neurons were more broadly tuned to temporally-modulated tone sequences (75 ms, p=0.009; 375 ms, p=0.04; 575 ms: p=0.004; Mann-Whitney test). Error bars, median±interquartile range.

**Extended Data Figure 8.**
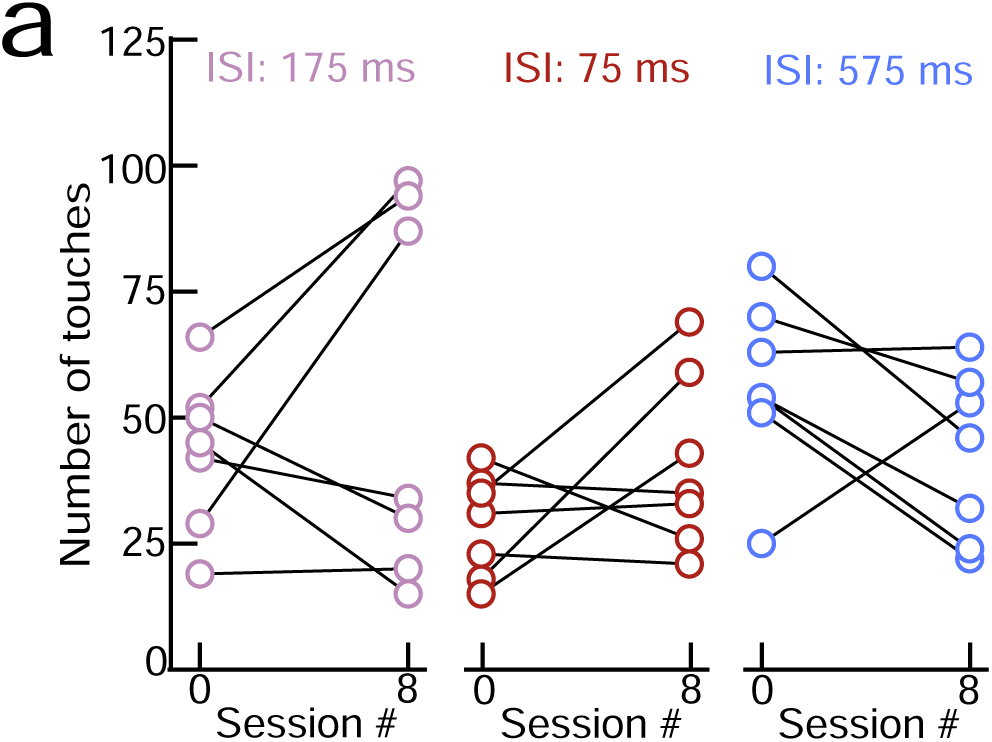
Pup-naive virgins did not increase the number of times they pressed a lever to turn off prototypes or morphs. **a,** The total number of levers presses in a session (which turns off continuously playing prototypes or morphs) did not increase by session 8, regardless of the stimulus group (N=7 mice per groups; S_0_ vs. S_8_: 175 ms, p=0.44, 75 ms, p=0.19, 575 ms, p=0.14; Student’s paired two-tailed t-test).

**Extended Data Figure 9.**
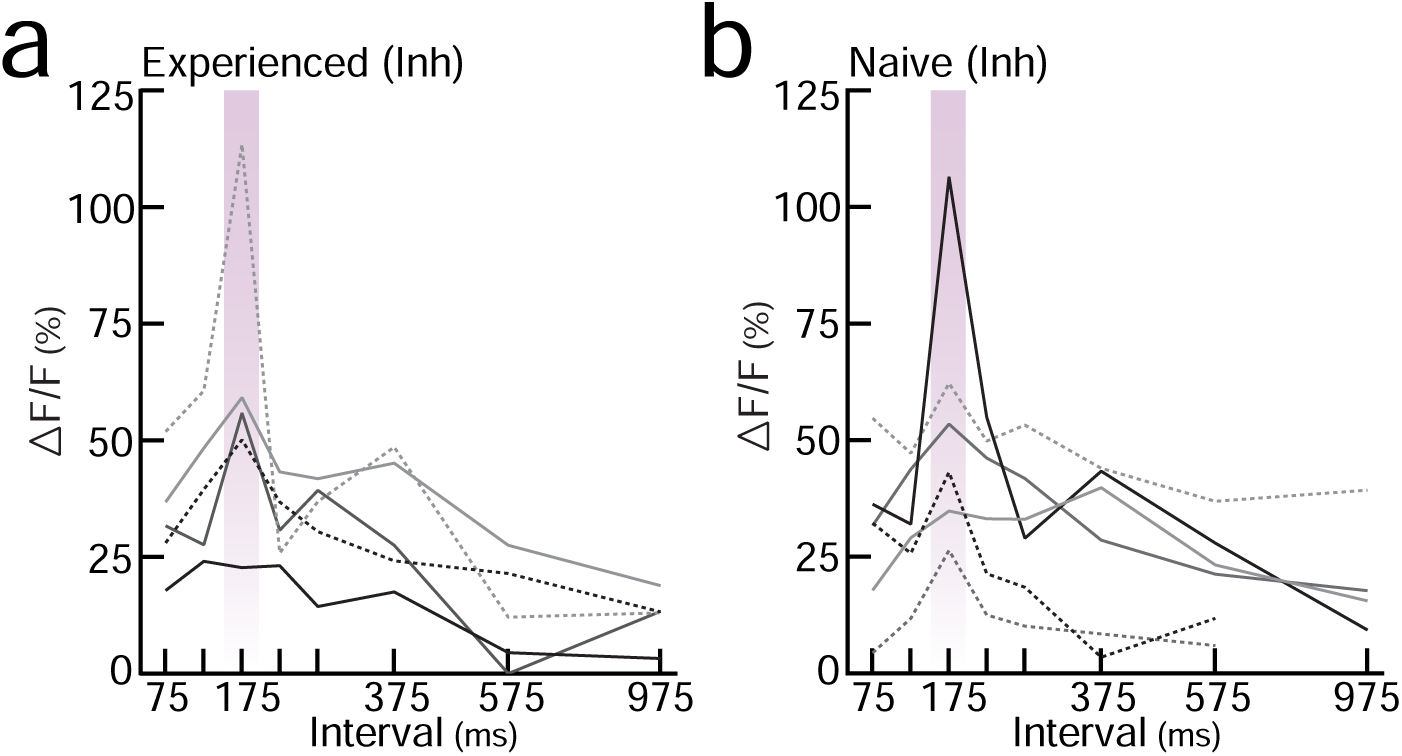
Temporal tuning curves for inhibitory neurons in experienced and naive auditory cortex. **a,** Inhibitory temporal tuning in experienced virgins (N=5 mice). Each line denotes an individual animal’s tuning curve. **b,** Inhibitory temporal tuning in naive virgins (N=6 mice).

**Extended Data Figure 10.**
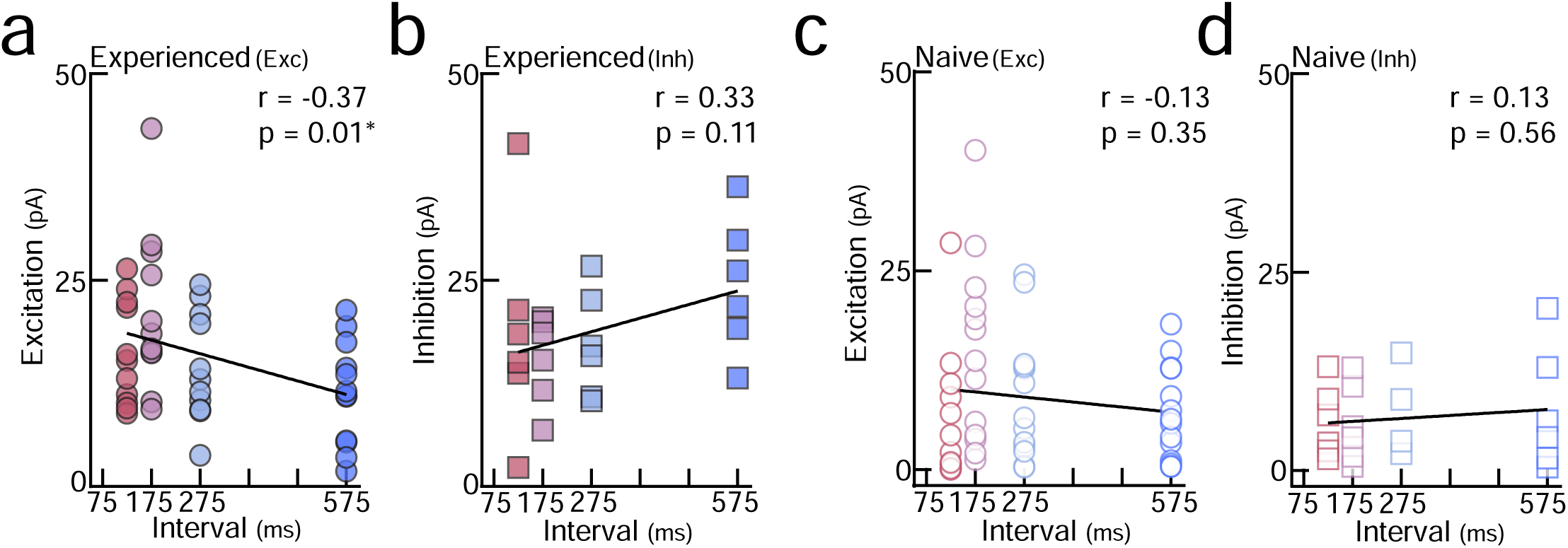
Evoked EPSCs track ISI duration in experienced cortex. **a,** Evoked EPSCs were negatively correlated with ISI duration in experienced females, with the weakest EPSCs evoked by slow morphs ∼575 ms (n=12 neurons; Pearson’s r=-0.37, p = 0.01). PSC (pA) = Area_(PSC)_ divided by stimulus duration (ms). **b,** Evoked IPSCs were not significantly correlated with ISI duration in experienced females (n=6; Pearson’s r=0.33, p=0.11). **c,d,** Evoked EPSCs (n=13) and IPSCs (n=6) were not correlated with ISI duration in naive cortex (EPSCs: Pearson’s r=-0.13, p=0.35; IPSCs: Pearson’s r=0.13, p=0.56).

**Extended Data Figure 11.**
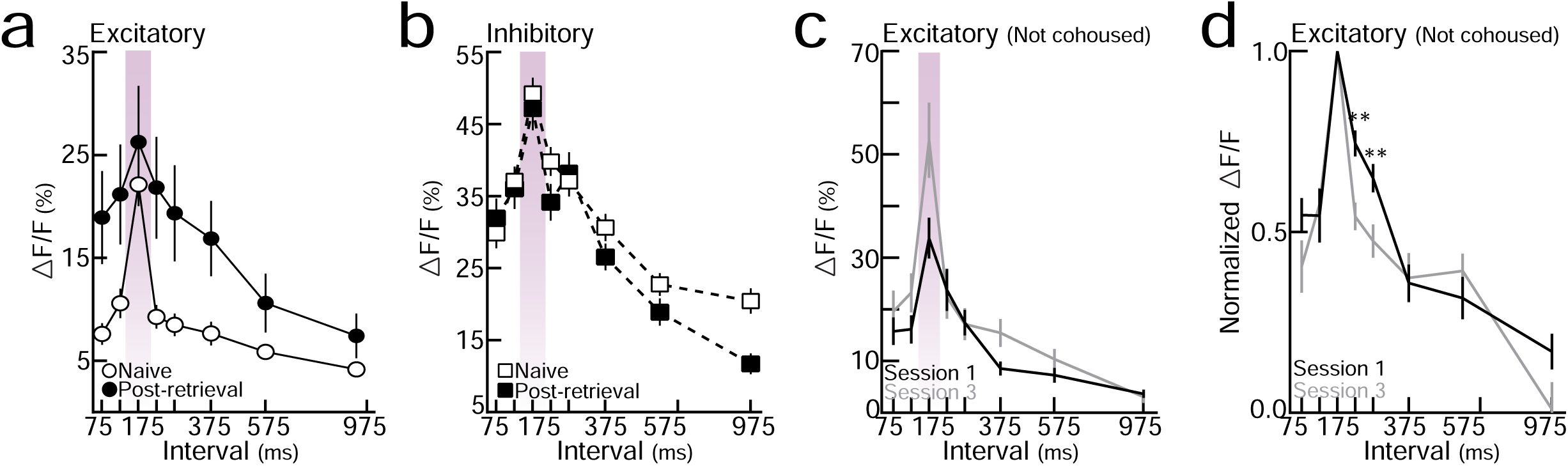
Responses in naive virgins are stable in absence of experience with pups. **a,** Temporal tuning across excitatory neurons before and after retrieval onset (raw data for Fig. 3h). **b,** Temporal tuning across inhibitory neurons before and after retrieval onset (raw data for Fig. 3h). **c,d,** To ensure that listening to pup calls under the two-photon could not explain the broadening of excitatory tuning we observed in Fig. 3, we assessed temporal tuning to pup calls in pup-naive virgins for three consecutive imaging days without cohousing or retrieval testing. There were no systematic changes in cortical tuning across temporal morphs in the absence of experience with pups (Session 1 (N=4 mice, n=43-52 single-cell tuning curves) vs. session 3 (N=5, n=27-48): 225 ms, p=0.0003; 275 ms, p=0.007).

**Extended Data Figure 12.**
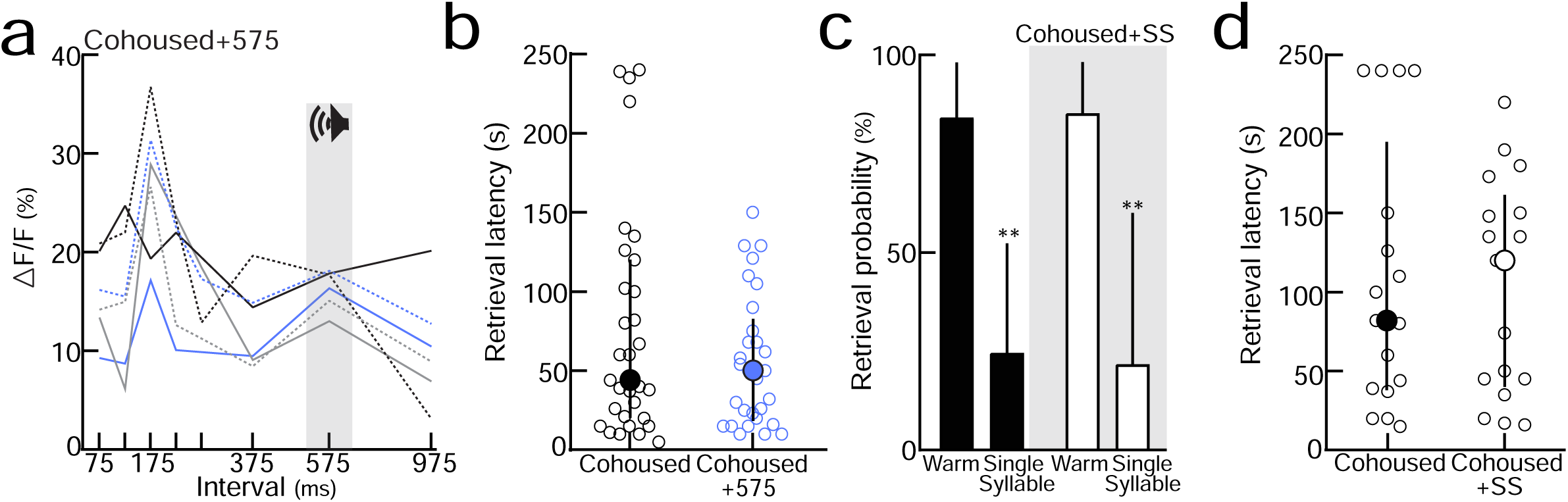
Playback of slow morphs during cohousing alters cortical tuning. **a,** Temporal tuning in virgins exposed to slow (575 ms) morphs throughout cohousing. Tuning assessed 24 hours after retrieval onset. Each colored line represents the tuning curve for one virgin. **b,** Latency to retrieve on successful trials from Fig. 4d (p=0.55; Mann-Whitney test). Median±Interquartile range. **c,** Playback of single syllable calls during cohousing did not alter the behavioral-salience of single syllables (‘Cohoused+SS’). Cohoused (N=6 mice): warm (n=11/13 trials) vs. cold-single syllable (n=2/9), p<0.001. Cohoused+SS (N=4 mice): warm (n=12/14 trials) vs (n=4/16), p<0.001; Binomial test. Errors bars, 95% binomial CI. **d,** Latency to retrieve on successful trials from **Extended Data Fig. 12c** (p=0.98, Mann-Whitney test). Error bars, median±interquartile range.

**Extended Data Figure 13.**
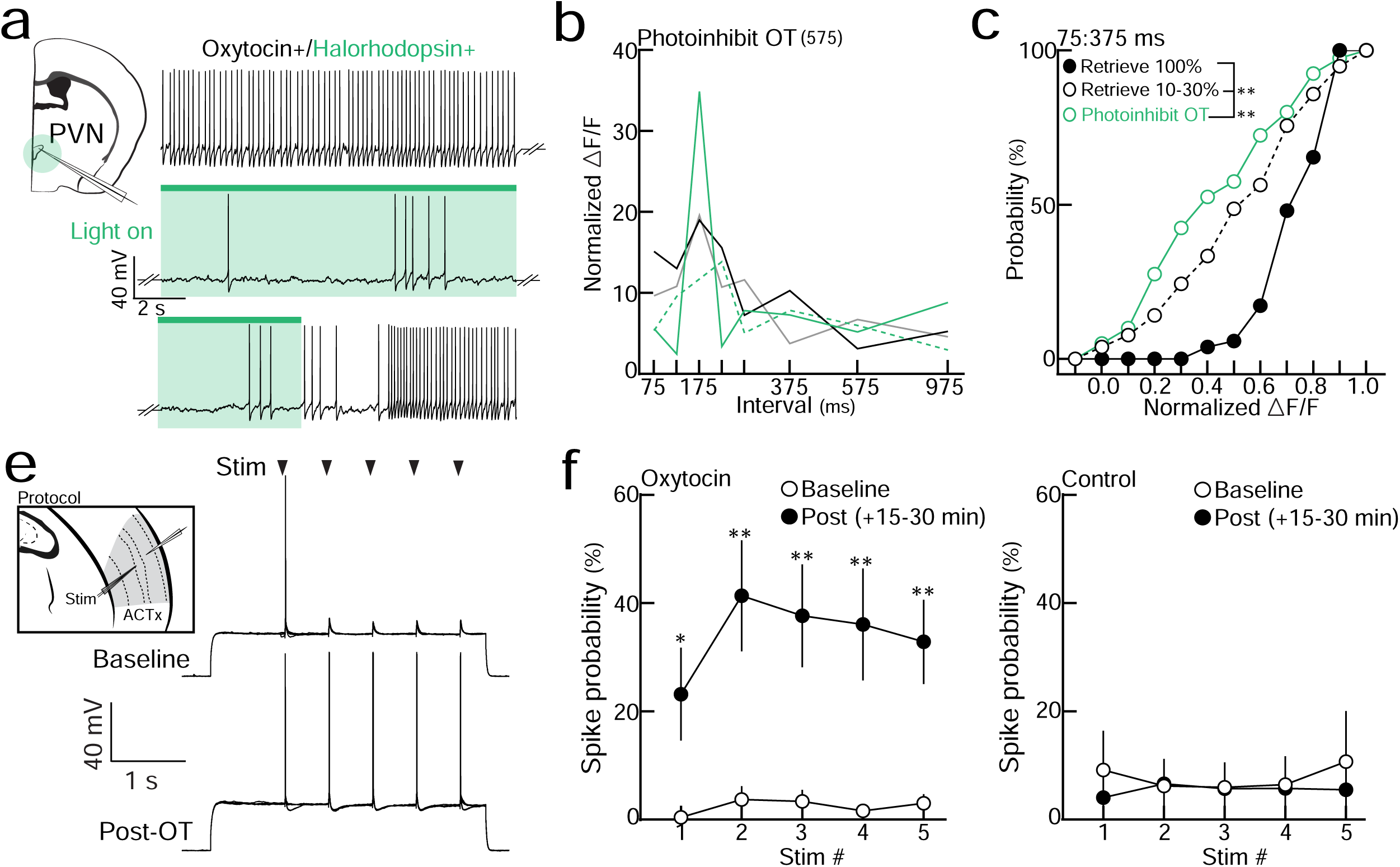
Oxytocin modulates cortical plasticity during cohousing. **a,** Example in vitro current-clamp recording from an oxytocin neuron containing the inhibitory opsin halorhodopsin. Green light efficiently perturbed spiking. **b,** Temporal tuning curves from cohoused virgins that received photoinhibition of OT neurons during 575 ms playback (‘photoinhibit OT_(575)_’). Each colored line represents the tuning curve for one virgin. **c,** Cumulative distributions of normalized ΔF/Fs averaged across behaviorally-salient ISIs (75:375 ms). Retrieve_100%_ vs. retrieve_10-30%_, p<0.0001, vs. photoinhibit OT_(575)_, p<0.0001; Kruskal-Wallis test. Retrieve_100%_ and retrieve_10-30%_ are same distributions as **Extended Data Fig. 6**. **d,** Example in vitro current-clamp recording from auditory cortex. The probability of evoking spikes in response to 5 extracellular stimulus pulses (ISI:575 ms) was measured before and after (1 µM) oxytocin wash. **e,** Evoked spike probability was significantly enhanced following oxytocin wash (post: 15-30 minutes; n=9 cells, For all stimulations, p<0.05). Repetitive stimulation in the absence of oxytocin (Control, n=7 cells) did not induce changes in spike probability (For all stimulations, p>0.05; Student’s paired two-tailed t-test).

## Acknowledgements

We thank I. Carcea, C.L. Ebbesen, W. Gan, E. Glennon, M. Insanally, K. Kuchibhotla, D. Lin, M.A. Long, N. López Caraballo, R. Oyama, and J.A. Schiavo for comments, discussions, and technical assistance. The AAV.mDLX.GcAMP6f virus **(Fig. 2a**) was a gift of J. Dimidschstein and G. Fishell. S.E. Ross created artwork in **Fig. 1b, 3a, 4b**, and **Extended Data Fig. 3**. We thank K. Furman and M. Hopkins for their help in developing the operant paradigm used in **Fig. 1i-k**. This work was funded by an NSF Graduate Research Fellowship (J.K.S. and K.A.M.); a Leon Levy Foundation Postdoctoral Fellowship and Brain & Behavior Research Foundation NARSAD Young Investigator Award (S.V.), as well as the BRAIN Initiative (NS107616), NICHD (HD088411), NIDCD (DC12557), a McKnight Scholarship, a Pew Scholarship, and a Howard Hughes Medical Institute Faculty Scholarship (R.C.F.).

## Author contributions

J.K.S conducted behavioral studies, in vivo optogenetics, and two-photon calcium imaging. S.V. performed in vivo whole-cell recordings. S.C.S and C.B.M performed in vitro whole-cell recordings. K.A.M wrote the code and made the hardware for operant testing. In vivo imaging and whole-cell data were analyzed by J.K.S; In vitro whole-cell data was analyzed by J.K.S, S.C.S, and C.B.M. J.K.S. and R.C.F designed the study and wrote the paper.

## References

1. Swain, J.E., Kim, P., Ho, S.S. Neuroendocrinology of parental response to baby-cry. J. Neuroendocrinol. 23, 1036–1041 (2011).

2. Lingle, S., Wyman, M.T., Kotrba, R., Teichroeb, L.J., Romanow, C.A. What makes a cry a cry? A review of infant distress vocalizations. Curr. Zool. 58, 698–726 (2012).

3. Dulac, C., O’Connell, L.A., Wu, Z. Neural control of maternal and paternal behaviors. Science 345, 765–770 (2014).

4. Lester, B.M. & Boukydis, C.F.Z. Infant Crying: Theoretical and Research Perspectives. (Plenum Press, New York, 1985).

5. Ehret, G., Koch, M., Haack, B., Markl, H. Sex and parental experience determine the onset of an instinctive behavior in mice. Naturwissenschaften 74, 47 (1987),

6. Krishnan, K., Lau, B.Y., Ewall, G., Huang, Z.J., Shea, S.D. MECP2 regulates cortical plasticity underlying a learned behaviour in adult female mice. Nat. Commun. 8 (2017).

7. Elyada, Y. M. & Mizrahi, A. Becoming a mother — circuit plasticity underlying maternal behavior. Curr. Opin. Neurobiol. 35, 49–56 (2015).

8. Marlin, B.J., Mitre, M., D’amour, J.A., Chao, M.V., Froemke, R.C. Oxytocin enables maternal behavior by balancing cortical inhibition. Nature 520, 499–504 (2015).

9. Ehret, G. Infant rodent ultrasounds – A gate to the understanding of sound communication. Behav. Genet. 35, 19–29 (2005).

10. Liu, R.C., Miller, K.D., Merzenich, M.M., Schreiner, C.E. Acoustic variability and distinguishability among mouse ultrasound vocalizations. J. Acoust. Soc. Am. 114, 3412–3422 (2003).

11. Lindová, J., Špinka, M., Nováková, L. Decoding of baby calls: Can adult humans identify the eliciting situation from emotional vocalizations of preverbal infants? PLoS One 10, e0124317 (2015).

12. Thiessen, E.D. What’s statistical about learning? Insights from modeling statistical learning as a set of memory processes. Philos. Trans. R. Soc. B Biol. Sci. 372 (2017).

13. Holt, L.L. & Lotto, A.J. Speech perception as categorization. Atten. Percept. Psycho. 72, 1218–1227 (2010).

14. Bizley, J.K & Cohen, Y.E. The what, where and how of auditory-object perception. Nat. Rev. Neurosci. 14, 693–707 (2014).

15. Petkov, C.I. & Jarvis, E.D. Birds, primates, and spoken language origins: behavioral phenotypes and neurobiological substrates. Front. Evol. Neurosci. 4 (2012).

16. Castellucci, G.A., Calbick, D., McCormick, D.A. The temporal organization of mouse ultrasonic vocalizations. PLoS One 13, e0199929 (2018).

17. Ehret, G. & Bernecker, C. Low-frequency sound communication by mouse pups (Mus musculus): wriggling calls release maternal behavior. Animal Behav. 34, 821–830 (1986).

18. Ehret, G. Left hemisphere advantage in the mouse brain for recognizing ultrasonic communication calls. Nature 325, 249–251 (1987).

19. Liu, R.C., Linden, J.F., Schreiner, C.E. Improved cortical entrainment to infant communication calls in mothers compared with virgin mice. Eur. J. Neurosci. 23 (2006).

20. Kuchibhotla K.V. et al. Parallel processing by cortical inhibition enables context-dependent behavior. Nat. Neurosci. 20, 62–71 (2017).

21. Hickok, G. & Poeppel, D. The cortical organization of speech processing. Nat. Rev. Neurosci. 8, 393–402 (2007).

22. Ocklenburg, S., Ströckens, F., Güntürkün, O. Lateralisation of conspecific vocalisation in non-human vertebrates. Laterality 18, 1–31 (2013).

23. Kim, H. & Bao, S. Selective increase in representations of sounds repeated at an ethological rate. J. Neurosci. 22, 5163–5169 (2009).

24. Dimidschstein, J. et al. A viral strategy for targeting and manipulating interneurons across vertebrate species. Nat. Neurosci. 19, 1743–1749 (2016).

25. Butts, D.A. & Goldman, M.S. Tuning curves, neuronal variability, and sensory coding. PLoS Biol. 4 (2006).

26. Kerlin, A.M., Andermann, M.L., Berezovskii, V.K., Reid, R.C. Broadly tuned response properties of diverse inhibitory neuron subtypes in mouse visual cortex. Neuron 67, 858–871 (2012).

27. Katlowitz, K.A., Picardo, M.A., Long, M.A. Stable Sequential Activity Underlying the Maintenance of a Precisely Executed Skilled Behavior. Neuron 98, 1133–1140 (2018).

28. Valtcheva, S. & Froemke, R.C. Neuromodulation of maternal circuits by oxytocin. Cell Tissue Res. 375, 57–68 (2019).

29. Mitre M. et al. A Distributed Network for Social Cognition Enriched for Oxytocin Receptors. J. Neurosci. 36, 2517–35 (2016).

